# Echo characterizes the desynchronization of gene expression and chromatin accessibility during cell-state transitions

**DOI:** 10.64898/2026.05.20.726634

**Authors:** Connor Finkbeiner, Dominik Otto, Manu Setty

## Abstract

Cell-state transitions during differentiation and disease involve coordinated changes across gene expression and chromatin accessibility, but these modalities do not change in lockstep. For example, regulatory elements can be primed before their target genes are expressed or remain accessible after expression ceases. This desynchronization between changes in gene expression and chromatin accessibility can manifest at the level of cell states. Understanding the drivers of this desynchronization can give insights into the molecular mechanisms underlying cell-state progression.

Here we introduce Echo, a statistical framework that identifies desynchronized cell states and the associated genes and regulatory elements from paired single-cell RNA and ATAC data. Echo States estimates cell-state density independently for each modality and compares them to determine which states are better resolved in RNA or ATAC. Echo Features then predicts feature values over each state space to identify the genes, regulatory loci, and transcription-factor motifs driving this desynchronization.

Applying Echo to the developing human fetal retina, we find that desynchronization is pervasive across every major cell population. Expression of cell-cycle genes resolves multipotent progenitors in gene expression but not chromatin accessibility, while fate priming resolves cycling neurogenic precursors in chromatin accessibility before gene expression. By combining desynchronized states and features along the cone trajectory, we reconstructed the regulatory logic of cone fate specification from multipotent progenitors, revealing a tight coupling between multipotency exit, cell cycle and lineage specification. Applying Echo to human hematopoiesis, we identified that the balance between stem-cell quiescence and differentiation is resolved more strongly in chromatin accessibility than in gene expression. Our results establish desynchronization as a pervasive, structured feature of differentiating systems, and Echo as a framework for characterizing the interplay between gene expression and chromatin accessibility during cell-state transitions.

## Introduction

Cell-state transitions during differentiation and disease involve coordinated changes across gene expression, chromatin accessibility, and other molecular modalities. While these modalities are tightly linked through regulatory mechanisms, changes in one modality are not immediately reflected in others, leading to transient desynchronization between them. This phenomenon has been characterized most extensively for chromatin accessibility and gene expression. For example, enhancers of lineage-specifying genes have been shown to be primed, opening prior to expression of their target genes across biological systems^1,2^ (**Fig. 1A**). Similarly, regulatory elements can remain accessible after expression of their target gene is shut down, have been observed to be accessible in lineages that do not express their target genes, or are shut down before their targets are ever expressed indicating chromatin plasticity^3,4^ **(Fig. 1A**).

**Figure 1:**
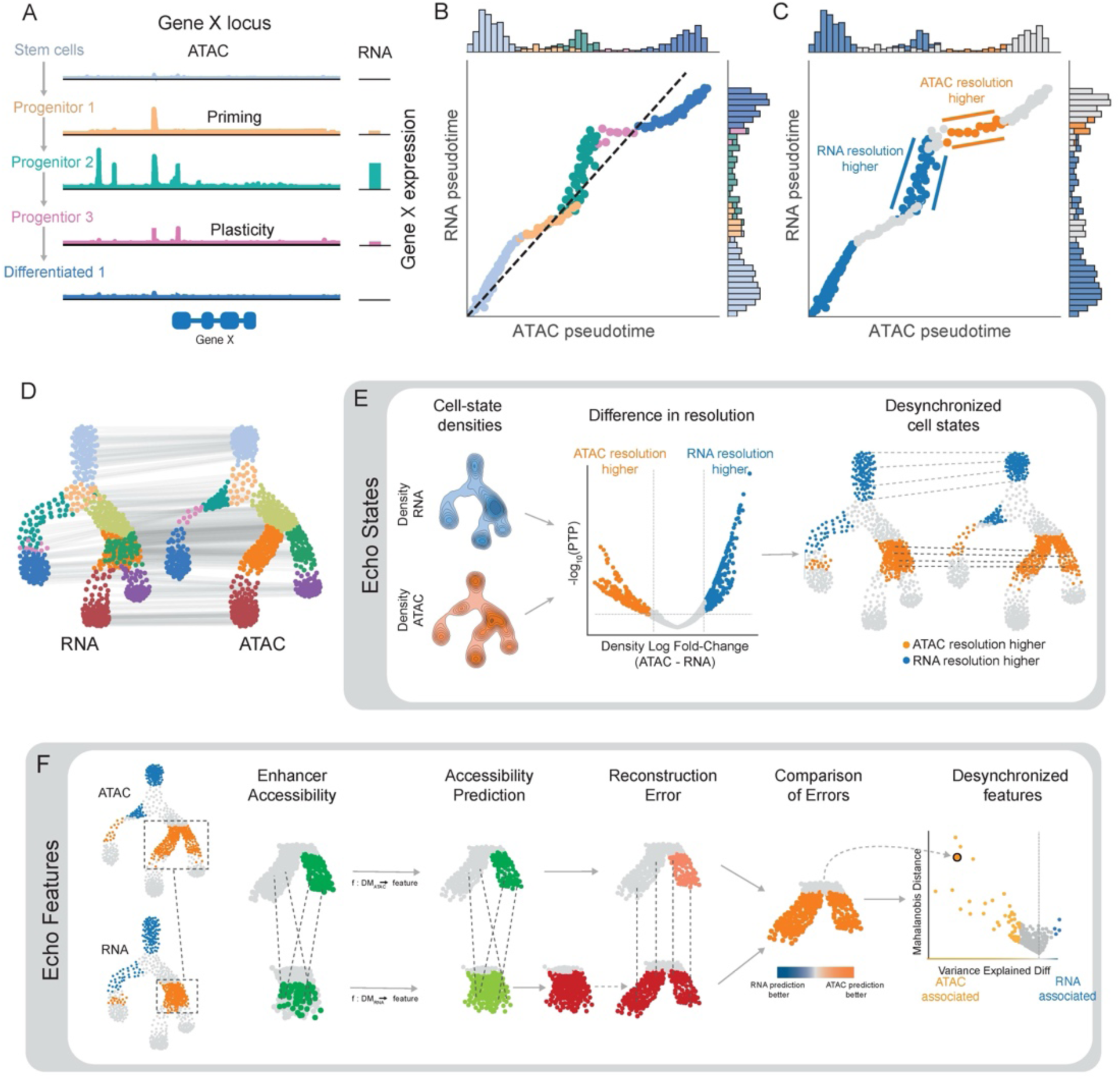
Echo modeling approach for identifying desynchronized states and features from paired single-cell RNA and ATAC-seq data. (A) Schematic of locus-level desynchronization between gene expression and chromatin accessibility during differentiation. Priming refers to gene loci that become accessible prior to gene expression; plasticity refers to loci that remain accessible after gene expression is shut down. (B) Locus-level desynchronization can be coordinated across multiple loci, producing desynchronization at the level of cell states. Schematic of theoretical pseudotemporal ordering of differentiating cells from (A): the RNA and ATAC pseudotime of the same cells do not progress in lockstep. Marginal histograms illustrate how cell densities along pseudotime differ between modalities. (C) Same as (B), highlighting cell states with significant resolution differences between RNA and ATAC modalities. (D) Schematic of paired RNA and ATAC state-space representations used as input to Echo. Lines connect the RNA and ATAC states of the same cells. (E) **Echo States**. Echo States leverages the inverse relationship between cell-state density and resolution to identify states better resolved in one modality than the other. Density functions are computed independently for each modality, and per-state density log fold-change quantifies the degree of desynchronization. Density uncertainties are used to compute a posterior tail probability (PTP) for significance. By convention, positive fold change indicates RNA-resolved states, and negative fold change indicates ATAC-resolved states. (F) **Echo Features**. Echo Features identifies the features underlying desynchronized states. The case of ATAC-resolved states with gene-level enhancer accessibility as the feature is shown. For each gene, enhancer accessibility is predicted as a function of the RNA and ATAC state spaces independently, and the difference in prediction accuracy is used to quantify the feature’s association with desynchronization. Mahalanobis distance provides a measure of statistical divergence between predictions from the two state spaces. The same approach applies to RNA-resolved states using gene expression as the feature. By convention, positive values indicate stronger association with RNA-resolved states, and negative values indicate stronger association with ATAC-resolved states.

The locus-level observations of priming and plasticity indicate that chromatin accessibility and gene expression do not change in concert but are instead temporally disconnected or desynchronized during cell-state transitions^5^. When such desynchronization is coordinated across multiple loci, it manifests at the level of cell states **(Fig. 1B, C**). Indeed, studies have observed primed cell states where regulatory loci of several genes are accessible due to binding of lineage regulators without an immediate upregulation of the target genes^6–8^. In this case, the chromatin accessibility of the cell is preparing for future gene expression changes and as a result, the state of the cell is better resolved in chromatin accessibility rather than gene expression. Similarly, acute signaling response is accompanied by massive transcriptional changes without equivalent changes in accessibility^9,10^ and processes like cell cycle where oscillation of genes across phases is driven by protein availability while the chromatin remains stable^10^, represent instances where the state of a cell is better resolved in gene expression rather than chromatin accessibility.

Therefore, across diverse biological processes, changes in expression and chromatin accessibility do not occur in lockstep (**Fig. 1B, C**), with one often leading or lagging the other suggesting that cell-state desynchronization is a pervasive property of differentiating systems. Determining which cell states are desynchronized and identifying the genes and regulatory elements that define this desynchronization, can inform the regulatory logic of cell-state transitions in differentiation and disease.

Multimodal single-cell technologies that measure paired RNA and ATAC in individual cells^11–13^ offer the opportunity to characterize desynchronization but there have been limited attempts to quantify it directly. Several tools have been developed to model the dynamics of cell-state change across modalities, but they either rely on correlation between expression and accessibility which does not account for desynchronization^1,11,14^ or model desynchronization only at individual genes^14^. Other studies have attempted to address desynchronization in the context of generating unified cell state representations^15–19^. While these tools have been powerful for cell type identification and atlas construction, they are not designed to determine where cell states are desynchronized. Finally, approaches that decompose multimodal data into shared and modality-specific components have provided useful representations but are primarily designed to improve integration rather than to characterize the biological basis of modality-specific signals^17,20^.

Here we introduce Echo, a novel statistical framework for identifying desynchronized cell states and their associated genes and regulatory elements from paired single-cell RNA and ATAC data. Echo consists of two components, Echo States and Echo Features. Echo States first builds representations separately from RNA and ATAC modalities and estimates cell-state density on each^21^. Cell-state density is an indicator of change in differentiating systems: Lower density corresponds to greater change in cell state, and vice versa. Echo States thus compares density changes between RNA and ATAC to identify desynchronized states and the modality that is better resolved. Echo Features then identifies the genes and regulatory elements driving this desynchronization. For states better resolved in RNA, Echo Features identifies the genes whose expression leads to higher resolution in RNA. Conversely, for states better resolved in ATAC, Echo Features identifies the regulatory loci and TF motifs associated with higher resolution in ATAC. Together, Echo provides a comprehensive framework for characterizing the interplay between gene expression and chromatin accessibility during cell-state transitions.

We applied Echo to characterize fetal retinal development and determined that every major cell population shows extensive desynchronization, with states better resolved in either RNA or ATAC. Cell cycle regulation, exit from multipotency, and fate specification were each associated with desynchronized states, and Echo identified the genes, regulatory loci, and motifs underlying each. By combining desynchronized states and features along the cone trajectory, we reconstructed the regulatory logic of cone fate specification from multipotent progenitors, revealing a tight coupling between multipotency exit, cell cycle and lineage specification. We further applied Echo to hematopoietic differentiation and identified that the balance between quiescence and differentiation in stem cells is better resolved in ATAC than in RNA. Our results suggest that desynchronization is a pervasive feature of cell-state transitions across biological systems. Echo is available as an open-source tool at github.com/settylab/scEcho.

## Results

### Echo modeling approach

During differentiation and disease, cells traverse through a series of states that collectively constitute the cell-state space. The position of a cell in this space is determined by its molecular profile i.e., gene expression in RNA state space and chromatin accessibility in the ATAC state space. In regions where cells are actively changing, they spread out across the state space, resulting in lower density. Since these fast-changing cells occupy well separated states, our ability to resolve differences between them is higher. In contrast, cells in a stable or slowly changing state accumulate at higher density and our ability to resolve differences between them is limited. Thus, there is an inverse relationship between cell-state density and resolution. Since RNA and ATAC capture different molecular features of the same cells, the state occupied by the same cell can be at different densities in the two modalities. Therefore, the states can be resolved differently by the two modalities: states with higher density in RNA than ATAC indicate greater change in ATAC than RNA and thus are better resolved in ATAC. Conversely, states with higher density in ATAC than RNA are better resolved in RNA.

Echo States leverages this inverse relationship to identify desynchronized cell states using paired single-cell RNA and ATAC data. Echo takes separate cell-state representations for RNA and ATAC as input (**Fig. 1D**) and computes cell-state density functions independently for each modality (**Fig. 1E**) using Mellon^21^. Mellon uses Gaussian Processes (GPs) to generate robust single-cell density estimates by sharing information across related states. Further, as GPs do not assume specific functional forms^22^ Mellon can estimate densities within complex single-cell landscapes^21^. Echo then computes a fold change of RNA and ATAC densities at each state, quantifying the degree of desynchronization (**Fig. 1E, Methods**). By convention, Echo fold changes are computed using the RNA modality as reference. A negative fold change therefore represents higher density in RNA and hence greater resolution in ATAC. Conversely, a positive fold change represents greater resolution in RNA. Density uncertainties are then used to compute a posterior tail probability (PTP) to assess whether the difference between densities is significant. States are thus classified as higher resolution in ATAC, higher resolution in RNA, or not significantly different (**Fig. 1E, Methods**).

Desynchronized states are a consequence of coordinated changes in molecular profiles of one modality without comparable changes in the other. For states better resolved in RNA, coordinated changes in expression across multiple genes drive these state changes without comparable changes in the chromatin landscape. Conversely, states better resolved in ATAC are driven by coordinated changes in accessibility across multiple enhancers without comparable changes in gene expression programs. Therefore, features which drive higher resolution in one modality must track with state changes in that modality, while varying less across the matched states in the other. As a result, when feature values are predicted from the cell-state space of each modality, desynchronized features are better predicted in the higher-resolution modality than in the lower-resolution one.

Echo Features uses this intuition to determine desynchronized features underlying desynchronized states. Echo uses Gaussian Process (GP) regression to predict each feature (gene expression for RNA, enhancer accessibility for ATAC) as a function of the RNA and ATAC state spaces independently (**Fig. 1F**). GPs produce robust predictions without assuming specific functional forms, accounting for noise in measurements and non-linear relationships between states and features^22^. For each feature, Echo computes the prediction error from each state space. Prediction error will be low when the feature variance is well described by the state space and high when feature variance is not associated with variance of the state space. Echo therefore compares the fraction of variance explained by each representation as a measure of feature desynchronization (**Fig. 1F**). By convention, Echo computes the difference as RNA minus ATAC: positive values indicate stronger association with RNA-resolved states and negative values indicate stronger association with ATAC-resolved states **(Fig. 1F, Methods**). An empirical null distribution is used to assess the significance of each feature’s association with desynchronization. Echo Features additionally computes a Mahalanobis Distance for each feature, which quantifies the statistical divergence between predictions from the two representations and can be used to rank features by their degree of desynchronization (**Fig. 1F, Methods**). For RNA-resolved states, gene expression is used as features whereas for ATAC-resolved states, gene enhancer accessibility or motif accessibility are used (**Methods**). The resulting desynchronized features can then be interpreted in the context of trajectory position, fate probabilities, and other cell-state properties for mechanistic interpretation.

### Desynchronization in the developing human fetal retina

To investigate desynchronization between gene expression and chromatin accessibility in the context of development, we applied Echo to a single-cell multiomic (ATAC+RNA) dataset of the early human fetal retina^23^ (**Fig. 2A**). Retinal neuronal differentiation is well studied with key lineage-specifying factors identified^24^, and aspects of their regulatory logic described^25–27^, making this a well-suited system to understand the biological basis of desynchronization.

**Figure 2:**
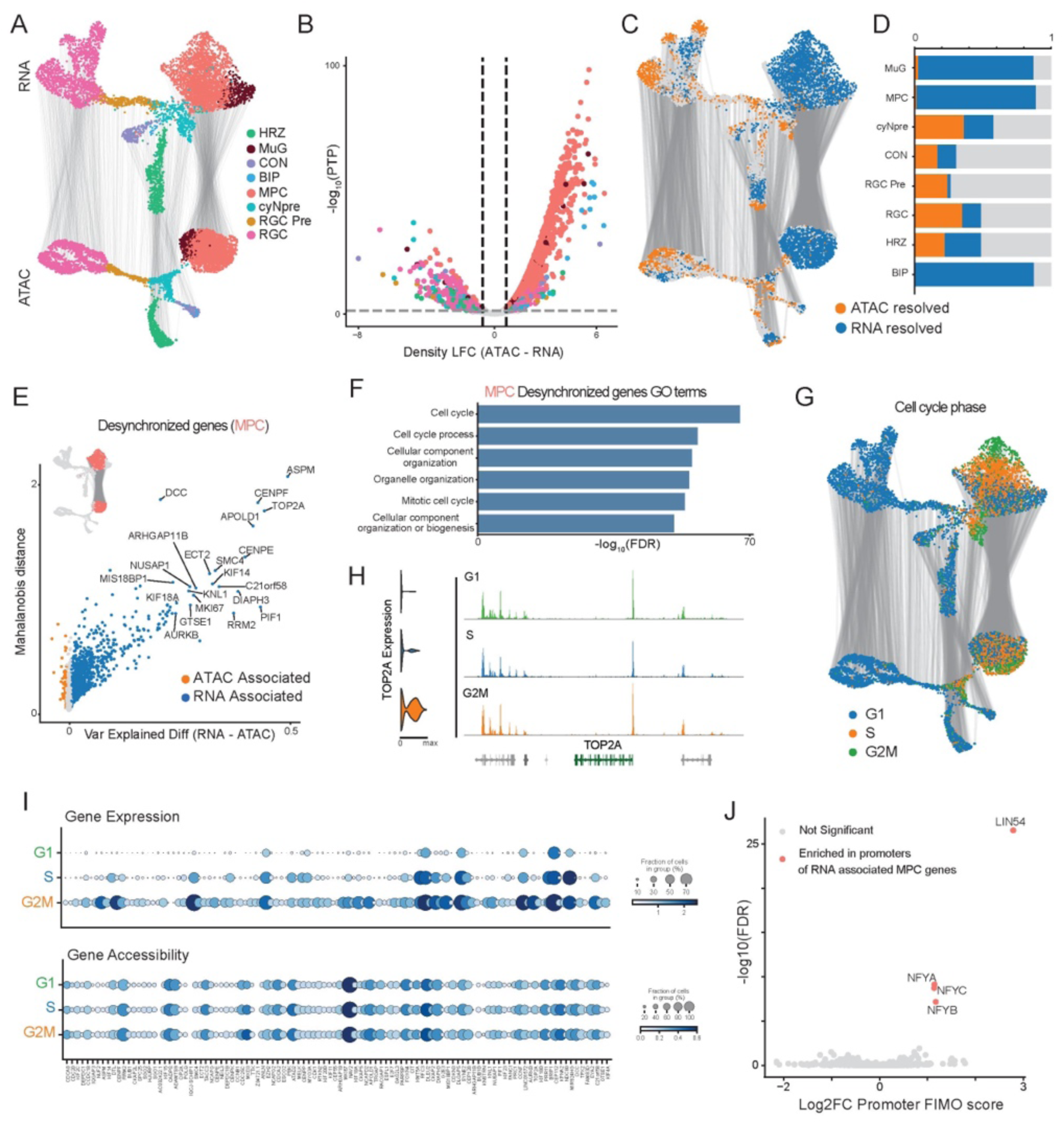
Desynchronization is pervasive in the developing fetal retina, with cell cycle resolving progenitor states in RNA but not ATAC. (A) UMAPs of Human Fetal retina D59 paired single-cell RNA- and ATAC-seq data^23^ colored by cell type. Top: RNA UMAP; bottom: ATAC UMAP. Lines connect the RNA and ATAC states of the same cells. MPC: Multipotent precursor cells; cyNpre: cycling neurogenic precursors; RGC: Radial glial cells; HRZ: Horizontal cells; MuG: Müller glia; CON; Cone cells; BIP: Bipolar cells. (B) Echo States volcano plot showing density log fold change between modalities (ATAC − RNA) on the x-axis and significance (posterior tail probability, PTP) on the y-axis. Cells passing thresholds for fold change and significance are colored by cell type. (C) Significantly desynchronized states from (B) projected onto the UMAPs from (A), colored by which modality resolves them better (ATAC-resolved in orange, RNA-resolved in blue). (D) Proportion of desynchronized states across all major cell types, illustrating that desynchronization is a pervasive feature of retinal development. (E) Echo Features applied to gene expression in the RNA-resolved MPC states. Pseudo-volcano plot shows difference in variance explained (RNA − ATAC) on the x-axis and Mahalanobis distance as the significance metric on the y-axis. Significantly desynchronized genes are colored by their modality association, and top-ranked genes are labeled. MPCs are highlighted in the inset on UMAPs from (A). (F) Gene Ontology terms enriched in the MPC desynchronized gene set, indicating that cell cycle drives RNA-resolved states in MPCs. (G) UMAPs from (A), colored by cell cycle phase. (H) Left: Expression of TOP2A (a top desynchronized MPC gene) across G1, S, and G2M phase MPCs from RNA. Right: Coverage plot showing accessibility of the TOP2A locus in the same cells from ATAC. (I) Top: Dot plot showing expression of all MPC desynchronized genes across G1, S, and G2M phase MPCs from RNA. Bottom: Dot plot showing accessibility of the same genes across the same cells from ATAC. Expression shows clear gradients across cell cycle phases, while accessibility remains comparable. nearly constant. (J) Promoter enrichment analysis of MPC desynchronized genes. X-axis: log fold change in mean promoter FIMO scores between the MPC desynchronized gene set and a background set of expression-matched genes. Y-axis: Mann-Whitney U test p-values adjusted by Benjamini-Hochberg correction. Significantly enriched motifs are highlighted.

Multipotent retinal progenitor cells (MPCs) generate all neuronal cells in the retina^28,29^. At this stage of development (Day 59), the differentiated cell types present are retinal ganglion cells, cone photoreceptors, and horizontal cells, along with a small number of early bipolar cells and Müller glia (**Fig. 2A**). As MPCs differentiate to neurons, they pass through a transitory state of cycling precursors expressing markers of the “last cell cycle” in progenitors, a state where fate decisions are thought to be made^30–33^. Since this transitory state was previously not annotated in this dataset, we reclustered the cells and identified a small group expressing these markers (**Supp. Fig. 1A-C**), which we labeled as cycling neurogenic precursors (cyNpre) (**Fig. 2A**).

We applied Echo States to identify desynchronized cell states, comparing cell state density across diffusion maps computed for the RNA and ATAC modalities separately (**Supp. Fig. 1D-E**). Across the retina, Echo States identified a variety of cell states with statistically significant differences in resolution between RNA and ATAC modalities (**Fig. 2B-D, Supp. Fig. 2A)**, indicating that desynchronization is pervasive in this system. Interestingly, while both the multipotent progenitors (MPCs) and the Müller glia, which have lost all differentiation potential, showed greater resolution in RNA, the differentiated neuronal cells and their precursors showed a mix of desynchronized states with greater resolution in either RNA or ATAC (**Fig. 2C-D**).

### Cell cycle resolves MPCs in RNA but not ATAC

MPCs have the highest proportion of desynchronized states among all major cell types, with a majority of these states are better resolved in RNA than ATAC (**Fig. 2D**). This pattern of desynchronization indicates that MPCs are undergoing widespread changes in gene expression without comparable changes in chromatin accessibility. We next applied Echo Features to identify these genes by comparing the predictive accuracy of gene expression between RNA and ATAC state spaces (**Fig. 2E)**. We observed that many of the top-ranking genes, including ASPM, TOP2A, and CENPE/F, are well-characterized cell cycle genes^34–36^. Further, gene ontology analysis of the full desynchronized gene set showed significant enrichment for cell cycle-related terms (**Fig. 2F**). These results indicate that cell cycle dynamics drive transcriptional changes in MPCs leading to greater resolution in RNA than in ATAC.

To test this directly, we annotated cells by their cell cycle phase^37^ (**Fig. 2G**) and compared expression and accessibility across them. The top desynchronized genes showed expression restricted to G2M MPCs, with little to no expression in other phases (**Fig. 2H, Supp. Fig. 2B**). Chromatin accessibility of these loci however remained near identical across all phases of the cell cycle (**Fig. 2H, Supp. Fig. 2C**). We observed the same pattern of G2M specific expression with comparable accessibility across all phases in all desynchronized genes (**Fig. 2I**). Further, this pattern was also maintained across all cell types and not just in MPCs (**Supp. Fig. 2D**). Thus, regulation of cell cycle genes in the retina appears to operate independently of chromatin accessibility, with changes in expression of cell cycle genes likely driven through changes in factor occupancy at already-accessible loci.

These results are consistent with studies that have identified cell cycle as one of the processes where gene expression and chromatin accessibility are decoupled^14^. Mechanistically, studies have suggested that expression of cell cycle genes is controlled through availability of transcription factors binding to constitutively open promoters in response to intracellular signaling^10,38^. We therefore compared transcription factor motif scores of promoters of the MPC desynchronized genes to a background set of non-desynchronized, expressed genes in MPCs. Our results showed significant enrichment for LIN54 (**Fig. 2J**, p < 1e-25, Wilcoxon rank sum test), a member of the DREAM complex, and the NFY factors (**Fig. 2J**, p < 1e-7, Wilcoxon rank sum test) both of which are known to regulate cell cycle genes^39,40^. Together, these results demonstrate that Echo can trace desynchronization from cell states to individual genes in cycling cells and uncover regulatory mechanisms governing this desynchronization.

### Fate priming leads to higher resolution in ATAC in cycling precursors

We next investigated the cyNpre population, the transitory cycling state where neuronal fate decisions are thought to occur^30–33^, since they showed the highest proportion of ATAC-resolved states in the developing retina (**Fig. 2D**).

Unlike MPCs, which showed higher resolution in RNA in the majority of states, the cyNpre cells showed a mixture of desynchronized states, with some cyNpre states better resolved in RNA and others better resolved in ATAC (**Fig. 3A**). We applied Echo Features and identified that cell states with higher RNA resolution in cyNpre are also driven by cell cycle genes similar to MPCs (**Supp. Fig. 3A-B**).

**Figure 3:**
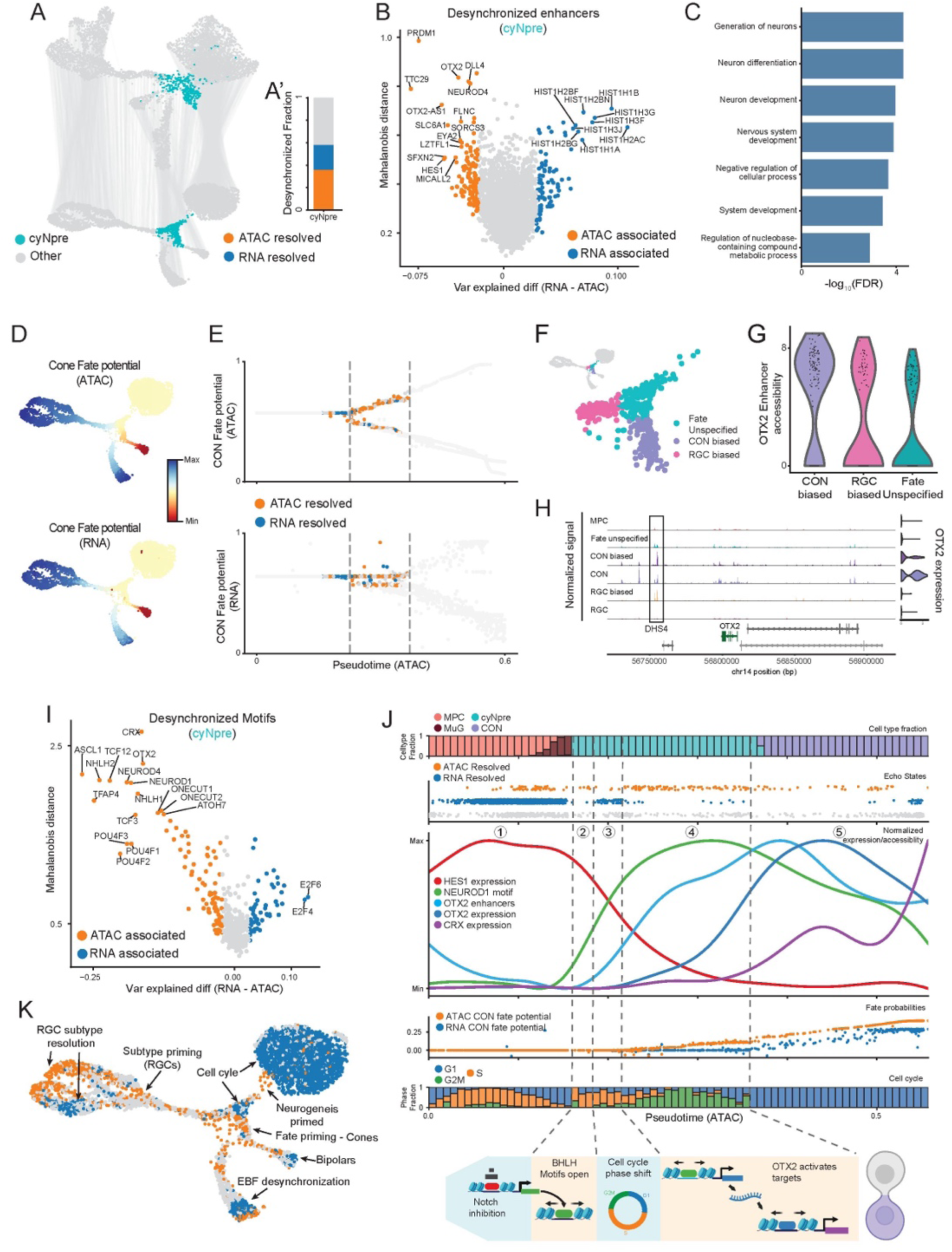
Cone fate trajectory is specified in ATAC prior to RNA in cycling neurogenic precursors. (A) UMAPs from Fig. 2A with cycling neurogenic precursors (cyNpre) highlighted. (A’) Bar plot shows the proportion of ATAC- and RNA-resolved states within cyNpre. (B) Echo Features applied to gene enhancer accessibility scores in the cyNpre cells. Pseudo-volcano plot shows difference in variance explained (RNA − ATAC) on the x-axis and Mahalanobis distance on the y-axis. Significantly desynchronized genes are colored by their modality association, and top-ranked genes are labeled. (C) Gene Ontology terms enriched in the cyNpre ATAC desynchronized gene set, indicating that loci associated with neurogenesis and fate specification are primed in cyNpre cells. (D) Top: ATAC UMAP from Fig. 2A colored by cone fate potential computed from ATAC using Palantir^47^. Bottom: Same as top, colored by cone fate potential computed using RNA. (E) Top: ATAC cone fate potential along ATAC pseudotime for all cells. Bottom: RNA cone fate potential along ATAC pseudotime. Desynchronized status of cyNpre status are shown, with all other cells as smaller gray dots. The highlighted window shows the pseudotime range in which cone fate is specified in ATAC prior to RNA. (F) cyNpre cells classified by fate bias based on ATAC cone fate potential from (D): cone-biased, RGC-biased, or fate-unspecified. Inset shows the same cells on the full ATAC UMAP. (G) OTX2 enhancer accessibility score (a top ATAC-resolved cyNpre desynchronized gene from (B)) across the cyNpre fate-bias groups from (F). (H) Coverage plots showing accessibility of OTX2 enhancers across cyNpre fate-bias groups from (F), with MPCs, mature cones, and RGCs included for reference. (I) Echo Features applied to chromVAR^74^ motif accessibility scores in cyNpre cells. Pseudo-volcano plot shows the difference in variance explained (RNA − ATAC) on the x-axis and Mahalanobis distance on the y-axis. Significantly desynchronized TFs are colored by their modality association, and top-ranked TFs are labeled. (J) Mechanistic interpretation of desynchronization along the cone fate trajectory. Top to bottom: cell type composition along ATAC pseudotime; Echo States classification (RNA-resolved, ATAC-resolved) along ATAC pseudotime; dynamics of key desynchronized features identified in (B) and (I), including HES1 expression, NEUROD1 motif accessibility, OTX2 enhancer accessibility, OTX2 expression, and CRX expression; ATAC and RNA cone fate potentials along ATAC pseudotime; and cell cycle phase composition along ATAC pseudotime. Bottom schematic: mechanistic interpretation of the trajectory, illustrating how multipotency exit and fate priming are coupled through the cell cycle. Dashed vertical lines mark changes in the modality of higher resolution along pseudotime.

We next sought to characterize the accessibility changes associated with higher ATAC resolution in the subset of cyNpre cell states. Since the relevant regulatory information may lie in enhancers rather than at the gene body alone, we constructed gene-level enhancer accessibility scores that aggregate accessibility across the correlated regulatory elements for each gene (**Supp. Fig. 4**, **Methods**). We then applied Echo Features to these enhancer scores, comparing how well the enhancer accessibility of each gene was predicted by the RNA and ATAC state spaces to identify regulatory loci whose accessibility tracks with ATAC-resolved states (**Fig. 3B**). Gene ontology analysis of this set revealed significant enrichment for generation of neurons, neuron development, and neuron differentiation (**Fig. 3C**), consistent with priming of neurogenic genes in cycling precursors. Notably, transcription factor OTX2, necessary for generation of both photoreceptor and bipolar cells^41–44^ and its target PRDM1, which regulates the fate decision between bipolars, and photoreceptors^45,46^ are among the top desynchronized genes.

Given that enhancers of broad neurogenic genes and specific regulators of cone fate are primed in cyNpre, we hypothesized that a subset of cyNpre cells are fate-primed and are biased towards the cone fate in ATAC prior to RNA. To test this, we computed fate probabilities independently from each modality using Palantir^47^ (**Fig. 3D**, **Supp. Fig. 5**). Cone fate probabilities increased in ATAC before RNA and this increase occurs specifically within the ATAC-resolved cyNpre states (**Fig. 3E, Supp. Fig. 5E**). These results imply that cone fate bias is established in chromatin within a subset of cyNpre cells, before this bias is reflected in gene expression.

We next sought to identify regulatory elements that are associated with cone fate specification in cyNpre cells. We first classified cyNpre cells as cone-specified, RGC-specified, or fate-unspecified based on their ATAC fate probabilities (**Fig. 3F**). Since OTX2 is known to be necessary for cone development^41,42^ and is one of the top desynchronized genes (**Fig. 3B**), we compared the enhancer accessibility scores of OTX2 across the three groups. OTX2 enhancer accessibility was highest in the cone-specified subset (**Fig**. **3G****, Supp. Fig. 6A**) and these enhancers were accessible prior to initiation of OTX2 expression (**Supp. Fig. 6B**). Amongst the enhancers assigned to OTX2, accessibility of the known OTX2 enhancer DHS4^48^ tracks with OTX2 enhancer accessibility score in fate-unspecified cyNpre cells, and increases in DHS4 accessibility precedes OTX2 gene expression, consistent with its role as the initiating enhancer^48^ (**Supp. Fig. 6C**). Therefore, by combining Echo results with fate probabilities and pseudotime ordering, we identified that cone fate is primed in a subset of cyNpre cells, with DHS4 accessibility as an important factor priming these cells toward cone fate prior to OTX2 upregulation (**Fig. 3H, Supp. Fig. 6D**).

Further, Echo Features also identified a smaller set of genes whose enhancer accessibility was better predicted by RNA states (**Fig. 3B**), implying that their accessibility tracks with RNA states better rather than ATAC states. This set was enriched for histone genes (**Supp. Fig. 3E**), whose expression is known to vary with cell cycle^49^, likely reflecting their coupling to RNA-resolved cell cycle dynamics.

### Regulatory logic of the cyNpre cells

Having identified the genes whose enhancer accessibility drives desynchronization, we next sought to identify the transcription factors driving these changes in chromatin accessibility by applying Echo Features to chromVAR motif accessibility scores in cycling neurogenic precursors (**Fig. 3I**). The desynchronized TFs fell into two broad categories: motifs for bHLH factors (ASCL1, NEUROD1, NEUROD4, ATOH7, BHLHE22) that are thought to drive MPCs to initiate differentiation^50–52^; and motifs for neuron fate-specifying factors including CRX^43,53–55^, OTX2^41,42^, the ONECUTs^41–43^, and POU4F2^56^. Thus, despite constituting less than 10% of cells, cyNpre cells show desynchronized features that are indicative of both exit of multipotency and fate specification. Given the known role of these cells as the primary intermediates between multipotent progenitors and differentiated neurons, we next investigated how desynchronized states and features unfold across differentiation progression. Specifically, we traced the desynchronization dynamics along the cone trajectory to characterize the temporal logic of regulatory events during specification and commitment of this lineage (**Fig. 3J**).

At the start of the trajectory, MPC states are better resolved in RNA, driven by cell cycle gene expression (**Fig. 2**, **Fig. 3J (1)**). The transition from MPCs into cyNpre, coincides with cells entering S-phase, and unlike MPCs is better resolved by ATAC (**Fig. 3J (2))**. This higher resolution in ATAC is likely driven by increased accessibility of bHLH motifs (**Supp. Fig. 7A**), implying that the early steps in neurogenesis are driven by chromatin remodeling before widespread upregulation of neurogenic gene expression. The increase in bHLH motif accessibility is coordinated with downregulation of HES1 (**Fig. 3J (2))**, consistent with the release of Notch-mediated repression that allows neurogenic bHLH factors to engage their target sites^52,57,58^. Cells then transition from S to G2M phase, with states now better resolved by RNA as cell cycle gene expression dominates (**Fig. 3J (3), Supp. Fig. 7B-C**). As cells exit the cell-cycle, desynchronization flips to better resolution in ATAC, driven in part due by increased accessibility at OTX2 enhancers and marked by a priming of cone fate potential (**Fig. 3J (4), Supp. Fig. 7C**). This includes accessibility of the DHS4 enhancer which is known to regulate OTX2 (**Supp. Fig. 7D**). DHS4 itself contains motifs for bHLH factors^48^, indicating a coupling of bHLH-driven exit from multipotency to OTX2-mediated fate specification through the cell cycle. OTX2 expression is subsequently upregulated, opening chromatin at new sites, though there is a delay before expression of its target genes such as CRX^41^. Eventual upregulation of these targets presumably commits cells to cone identity^41,53^ (**Fig. 3J (5**)).

Thus, Echo highlights that desynchronization plays an important role in differentiation and cell fate specification in the retina and identifies motifs, genes, and enhancers that are known to play critical roles in retinal differentiation as underlying these states. Further, the temporal ordering of features by their modality of change shows that exit from multipotency, cell cycle progression, and fate specification are tightly coupled revealing the regulatory logic by which these factors interact to confer cone fate.

More broadly, Echo characterized desynchronized states in every major population, with distinct regulatory mechanisms underlying each from cell cycle regulation in MPCs to resolution of subtypes in retinal ganglion cells (**Fig. 3K**). Together, these findings suggest that desynchronization as a pervasive and structured feature of retinal development, with the modality in which each state resolves reflecting the regulatory process that defines it.

### Desynchronization in Hematopoiesis

We next investigated whether desynchronization between expression and chromatin accessibility is a feature of other differentiating systems by applying Echo to a single-cell multimodal data of CD34+ hematopoietic stem and progenitor cells from human bone marrow^59^ **(Fig. 4A**). Hematopoiesis is a homeostatic differentiation system where hematopoietic stem cells (HSCs) continually produce blood and immune cells. Unlike retinal development, where the progenitor cells fully differentiate during development, hematopoietic differentiation maintains a pool of long-term quiescent hematopoietic stem cells (HSCs) that retain both renewal and differentiation abilities. We applied Echo States to hematopoietic differentiation and similar to retinal development, detected desynchronized cell states across all cell types (**Fig. 4B-C**).

**Figure 4:**
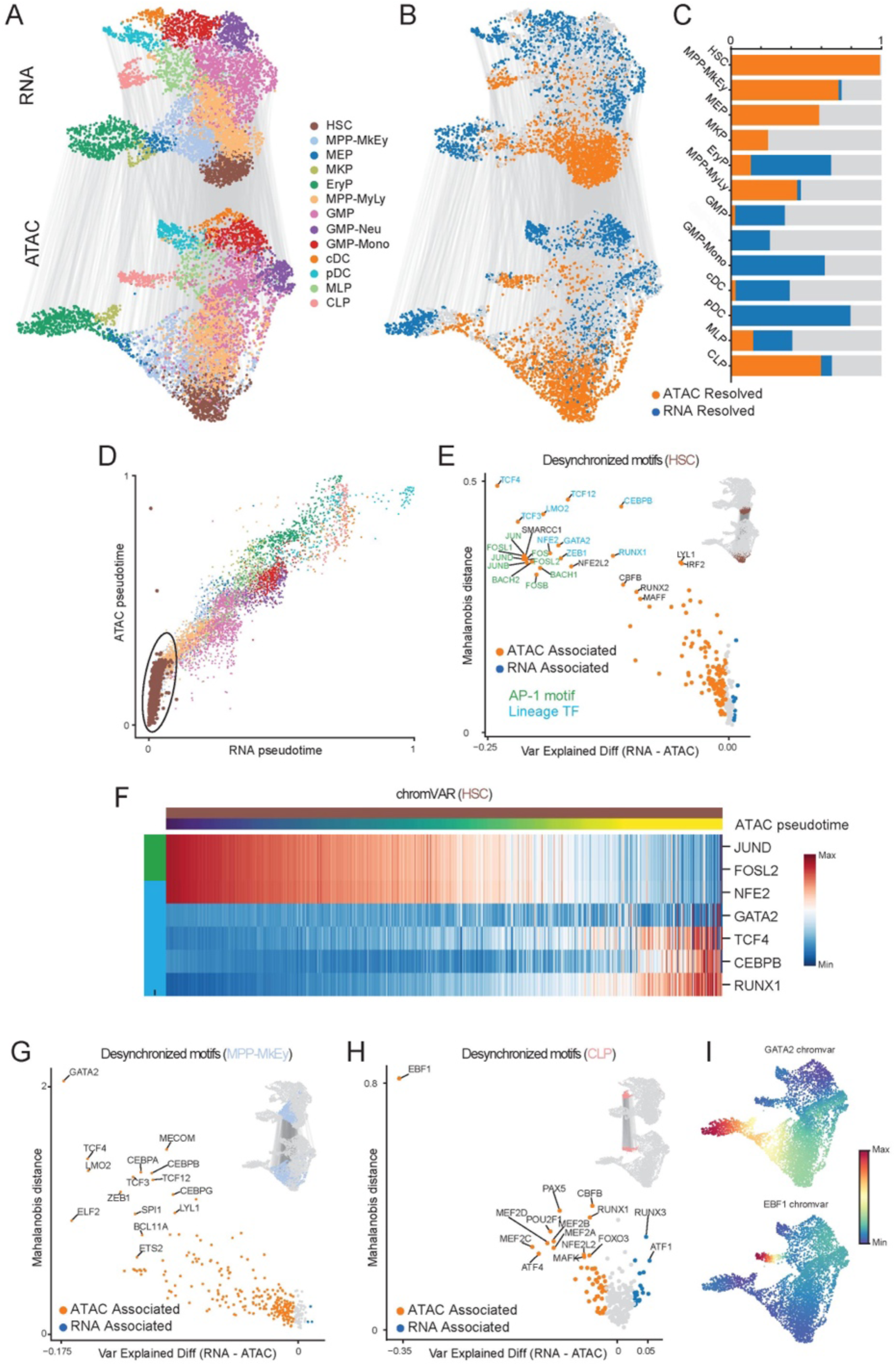
Desynchronization is pervasive in hematopoietic differentiation, with the quiescence-differentiation balance resolved in ATAC but not RNA. (A) UMAPs of CD34+ hematopoietic stem and progenitor cells from^59^ colored by cell type. Top: RNA UMAP; bottom: ATAC UMAP. Lines connect the RNA and ATAC states of the same cells. (B) UMAPs from (A), colored by Echo States results (ATAC-resolved in orange, RNA-resolved in blue). (C) Proportion of desynchronized states across all major cell types, illustrating that desynchronization is a pervasive feature of hematopoietic differentiation. (D) Scatter plot of ATAC pseudotime vs. RNA pseudotime for all cells, colored by cell type. HSCs (circled) show higher resolution in ATAC than RNA. (E) Echo Features applied to chromVAR motif accessibility scores in HSCs. Pseudo-volcano plot shows the difference in variance explained (RNA − ATAC) on the x-axis and Mahalanobis distance on the y-axis. Significantly desynchronized TFs are colored by their modality association, and top-ranked TFs are labeled. AP-1 motifs are labelled in green and hematopoietic lineage regulators are labelled in cyan. Inset: HSCs highlighted on UMAPs from (A). (F) Heatmap showing chromVAR motif accessibility scores for select AP-1 and hematopoietic lineage factors along ATAC pseudotime for HSCs. AP-1 factors show high accessibility at early stages and decline along HSC differentiation, while lineage regulators show the opposite pattern. (G) Echo Features applied to chromVAR motif accessibility scores in erythroid progenitors. Pseudo-volcano plot shows the difference in variance explained (RNA − ATAC) on the x-axis and Mahalanobis distance on the y-axis. Significantly desynchronized TFs are colored by their modality association, and top-ranked TFs are labeled. Inset: Erythroid lineage cells (MPP, MkEy) highlighted on UMAPs from (A). (H) Same as (G) for CLPs. (I) ATAC UMAPs colored by GATA2 (top) and EBF1 (bottom) chromVAR motif accessibility scores. GATA2 and EBF1 are the top ATAC-resolved desynchronized TFs from (G) and (H), respectively, and are master regulators of erythroid and lymphoid lineages.

We first characterized the RNA-resolved states in plasmacytoid dendritic cells (pDCs), as these cells showed the highest degree of RNA-driven desynchronization (**Fig. 4C**). We applied Echo Features to determine the genes driving this desynchronization and observed significant enrichment of cell cycle genes among this set (**Supp. Fig. 8A-B**). This is consistent observation made in retinal progenitor cells, where we also identified that expression of cell cycle genes is regulated independently of chromatin accessibility.

More broadly, hematopoietic cells enter and exit the cell cycle multiple times and cycling cells can be observed across all lineages (**Supp. Fig. 8C**). Therefore, we next examined whether cell cycle underlies all RNA-driven desynchronization in hematopoiesis (**Fig. 4C**). We observed that cells in G2M or S phases include the vast majority of cell states better resolved in RNA (**Supp. Fig. 8D**) and the majority of ATAC-resolved states are in G1 **(Supp. Fig. 8D**). Further, there is overlap in genes associated with higher resolution in RNA within cycling cells in different hematopoietic cell types (**Supp. Fig. 8E**), suggesting that cell cycle operates through a core set of genes regulated by a common post-translational mechanism. Together with the retinal findings, these results indicate that cell cycle-driven RNA desynchronization is a consistent feature across differentiating systems.

### ATAC provides better resolution into the factors controlling HSC quiescence and differentiation

HSCs had the highest proportion of ATAC resolved states in hematopoiesis with virtually all HSC states better resolved in ATAC compared to RNA (**Fig. 4C**). Interestingly, HSCs occupy a larger dynamic range in ATAC pseudotime with relatively smaller change in RNA pseudotime (**Fig. 4D, Supp. Fig. 9A**). This indicates that the earliest stages of hematopoietic differentiation, from long-term quiescent HSCs to differentiation-committed short-term HSCs^60^, are better resolved in chromatin accessibility than gene expression.

Since transcription factors are known to be critical regulators of hematopoietic differentiation^61^, we applied Echo Features to chromVAR motif accessibility scores to identify TFs associated with this ATAC-driven desynchronization in HSCs (**Fig. 4E**). The desynchronized TFs fall into two broad categories: hematopoietic lineage regulators such as TCFs, GATA2, NFE2, ZEB1, CEBPB, RUNX1 and LMO2^61,62^, and AP-1 family members such as JUN, FOS, and BACH factors (**Fig. 4E)**. Interestingly, these two groups show opposing patterns along the HSC trajectory: AP-1 motif accessibility decreases as HSCs progress toward differentiation, while lineage regulator accessibility increases (**Fig. 4F, Supp. Fig. 9B**).

To attribute functional roles to the AP-1 signal, we examined the expression patterns of individual AP-1 family members (**Supp. Fig. 4_2C**). We identified JUND, FOS, FOSL2, and FOSB as the likely drivers, based on their progressive downregulation along the HSC trajectory **(Supp. Fig. 9C**). Many of these factors have been implicated in HSC quiescence maintenance: sustained FOS expression locks HSCs in G0/G1 phase^63^, and FOSL2 loss leads to HSC expansion^64^. The opposing accessibility patterns of these quiescence-associated AP-1 factors and lineage regulators suggest a balance between quiescence maintenance and differentiation progression that is reflected more strongly in accessibility than in gene expression. The greater resolution in ATAC likely reflects this competition playing out across hundreds of accessible loci, whereas expression involves only a handful of genes.

Previous studies have demonstrated that ratio of expression between competing TFs can drive fate choice in hematopoiesis^47^. Our results suggest a potential mechanistic basis for such phenomenon in HSC quiescence where TFs may compete to regulate their respective targets and possibly to bind at the same loci. In support of this, lineage regulator NFE2 recognizes the same motif as AP-1 family members^65^. Further, Echo Features identified enrichment of signaling-related genes among the ATAC-driven desynchronized set (**Supp. Fig. 9D, E**), consistent with the known role of AP-1 factors as signal integrators and suggesting that the quiescence-differentiation balance is also modulated through signaling inputs. Together, these results indicate that ATAC-driven desynchronization in HSCs captures the balance between quiescence and differentiation commitment.

### Desynchronization in hematopoietic lineages

In addition to HSCs, we also observed significant ATAC-driven desynchronization in common lymphoid progenitors (CLPs) and megakaryocytes-erythroid progenitors (MPP-MkEy). We applied Echo Features using chromVAR scores and identified EBF1 and GATA2 to respectively be the strongest desynchronized TFs in these lineages (**Fig. 4G, H**). EBF1 is the master regulator of lymphoid lineage^66^ and GATA2 specifies megakaryocyte-erythroid lineages^67^, and thus the ATAC driven desynchronization led by these factors reflects the onset of the regulatory cascade triggered by these TFs for lineage specification. These results collectively demonstrate that desynchronization is a pervasive feature of differentiating systems, and that Echo can decompose it into distinct regulatory programs operating at different stages of the differentiation hierarchy.

## Discussion

Echo is a framework for comprehensively characterizing desynchronization between gene expression and chromatin accessibility from paired single-cell RNA and ATAC-seq data. Echo leverages the connection between cell-state change and cell-state density to identify states that are better resolved in one modality than the other. These resolution differences arise from coordinated changes in expression of multiple genes or accessibility at multiple regulatory loci without commensurate change in the other. Echo identifies these underlying features by comparing their predictive accuracy in each modality’s state space. This two-step framework transforms desynchronization from a phenomenon obscured by integration into an interpretable readout of the dynamic interaction between gene expression and chromatin accessibility as they guide cell-state transitions.

We applied Echo to characterize desynchronization during human fetal retina development and determined that desynchronization is a pervasive feature of every cell type. Multipotent progenitor cells showed RNA-resolved states driven by cell cycle, while cycling neurogenic precursors showed ATAC-resolved states driven by fate priming. Notably, individual cell types contained mixtures of both RNA- and ATAC-resolved states, pointing to a temporal interplay of expression and accessibility changes even within closely related cells, a heterogeneity that Echo’s single-cell resolution makes detectable. To interpret these patterns, we combined Echo results with trajectory descriptors including pseudotime and fate probability, reconstructing the regulatory logic that drives progenitors to cone photoreceptor cells. This revealed that progenitors lose their renewal potential simultaneously with fate priming, coupled through the cell cycle. Our work thus was able to recover known regulators of this process characterized over multiple studies, clarify the order of their functional interactions and suggest a novel link between cell cycle and fate specification in retinal development.

Our work also highlighted that hematopoietic differentiation, like retinal development, is punctuated by desynchronized states across all cell types. In addition to identifying desynchronization associated with known regulators of hematopoietic lineages, we identified that the balance between quiescence and differentiation in hematopoietic stem cells plays out in chromatin accessibility rather than gene expression, through a possible competition between AP-1 factors maintaining the quiescence program and lineage regulators driving differentiation. The findings across retina and hematopoiesis together highlight the significance of characterizing desynchronization to interpret differentiation trajectories. More broadly, paired single-cell data carry rich regulatory information across modalities, but realizing this potential has remained limited by the lack of approaches that resolve modalities rather than integrate them. Echo provides a framework to begin to unlock this information.

Desynchronization likely has implications for cross-modality imputation, where the goal is to reconstruct missing modalities using reference or bridge datasets^53^. These methods often rely on the assumption that the cell-state space of the missing modality can be reconstructed from the space of the measured modality. Our results suggest this assumption may not always hold, since the two modalities carry different state-level information in desynchronized states. Mapping from a lower-resolution to a higher-resolution modality therefore introduces uncertainties that need to be accounted for.

Two limitations of Echo should be noted. First, when concurrent but separate processes operate in both modalities, they may cancel at the state level, or only the dominant effect will be detected. Echo States therefore identifies relative resolution differences between modalities and cannot resolve cases where multiple modality-specific processes coexist at comparable strength. Similarly, if a single feature drives distinct processes in the two modalities, such as a transcription factor simultaneously upregulating one set of genes while priming the loci of others, and the net effects on each state space are equivalent, Echo will not detect this as desynchronization. The dominant effect, however, will still be identified. Second, our state-space construction is necessary for cross-modality density comparison but does not provide formal theoretical guarantees for this comparison. Empirically, however, across every dataset we tested, a subset of cell states consistently emerge with density fold change near zero. The biological identity of these states is not fixed across systems, but they reliably serve as a reference point: all Echo States results can be interpreted as deviation from these implicitly chosen reference states. More detailed empirical analyses and in silico perturbations to formally characterize these properties will be presented in a subsequent version of this manuscript.

## Methods

### Echo

The state of a cell is determined by its molecular profile. Coordinated changes in molecular profiles lead to changes in cell state, with the magnitude of state change correlated with the extent of profile changes. The state reflected in distinct molecular modalities such as gene expression and chromatin accessibility is not necessarily identical for any given cell, due to temporal offsets between them. Such temporal offsets can lead to desynchronization, when changes in one modality are not matched by commensurate changes in the other. Echo is designed to characterize such desynchronization between gene expression and chromatin accessibility. Using paired single-cell RNA- and ATAC-seq data as input, Echo identifies cell states where expression and accessibility are desynchronized, and the features (i.e. genes, regulatory elements, and transcription factors) driving this desynchronization. To achieve these objectives, Echo has two core components: Echo States and Echo Features. Echo States is a statistical framework for identification of desynchronized states and Echo Features is a framework for identification of genes and regulatory loci underlying desynchronized states. It is important to note that Echo is designed to detect relative differences between the two modalities. A finding of greater resolution or feature association in one modality indicates greater variation in that modality but does not imply that the other modality is stable.

#### Echo States

Cell-state space in our work refers to a low-dimensional embedding of cells based on their molecular profiles, in which proximity represents biological similarity. Fundamental biological processes such as proliferation, apoptosis, response to signaling, as well as molecular fluctuations shape how cells are distributed in this state space^21^. Specifically, regions that are traversed quickly are sparsely populated whereas slow changing cells lead to accumulation. Thus, how the cells are distributed in the state space, cell-state density, reflects the biological properties of the system and the measured modality^21^. Cell-state density also determines the resolution at which states can be distinguished: in sparse regions, cells are well separated and fine-grained differences between molecular states are recoverable, whereas in dense regions such differences are not similarly discernable. Density and resolution are thus inversely related. With paired RNA and ATAC measurements from single-cell multiomic data, each cell occupies a position in the RNA state space (derived from gene expression) and a separate position in the ATAC state space (derived from chromatin accessibility). Consequently, if one modality is changing faster than the other, cell-state density is likely to be lower in that modality, and the same population of cells will be resolved better there. This is the property Echo leverages to identify desynchronization.

Echo’s approach to identifying desynchronized cell states rests on two key assumptions. First, the cell-state space is constructed such that Euclidean distance between cells reflects biological similarity within each modality. This assumption is necessary for computation of cell-state density. Second, local density estimates derived from each modality are comparable across modalities. This assumption is necessary for quantifying the difference between densities.

##### Cell-state spaces

To satisfy these assumptions, Echo uses diffusion maps^68^ for cell-state space representations. Diffusion maps have been widely used for single-cell data since Euclidean distance in diffusion map space captures biological similarity more faithfully than distances in other representations^47,69^. Further, we have previously demonstrated that cell-state density in diffusion maps can represent biologically meaningful variability^21^, supporting its use as the basis for density estimation.

Diffusion maps are computed separately for the RNA and ATAC modalities from single-cell multiomic data after modality-specific preprocessing. Echo uses the diffusion maps implementation in Palantir^47^, which constructs the neighborhood graph using a fixed number of nearest neighbors and adaptive scaling. This construction renders density estimates invariant to the scale of the input feature space, a property relevant to the second assumption: Since RNA and ATAC are measured in very different feature spaces, scale invariance is necessary for their densities to be comparable.

Beyond scale invariance, we relied on empirical observations to assess whether densities are comparable across modalities in practice. We found that this construction reliably yields a density fold change of zero for a subset of cell-states across all examined datasets. This that would not arise if the two modalities were on systematically different scales, and we take that as evidence that comparability holds in practice.

##### Cell-state density

We use Mellon^21^ to estimate cell-state density from each modality. Mellon uses a Gaussian Process (GP) to infer a density function over the cell-state space. For each cell 𝑖 ∈ {1, …, 𝑛} and each modality 𝑘 ∈ {𝑅𝑁𝐴, 𝐴𝑇𝐴𝐶}, Mellon estimates a posterior mean log-density value 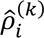 along with a posterior variance 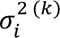 of the log-density estimate.

###### Intrinsic dimensionality

A practical requirement for cross-modality density comparison is that the two density estimates must be expressed in the same units. Mellon’s density estimator depends on the intrinsic dimensionality of the data, and density values computed under different dimensionalities are not directly comparable. To ensure that the RNA and ATAC density estimates are comparable, both density estimators are fit with a shared intrinsic dimensionality 𝑑^∗^, defined as:

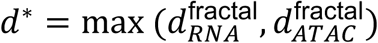

where 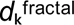 is the fractal intrinsic dimensionality of the embedded data of modality *k*, computed as described in Mellon^21^. In practice we found that 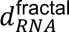 and 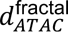 were typically close and as result, 𝑑^∗^ differs only modestly from either.

##### Comparison of densities across modalities

Desynchronization between modalities is quantified by the difference in log-densities between modalities:

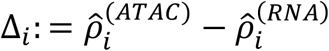

Δ*_i_*(𝑥) ≈ 0 represents states marked by their low density-difference, and greater deviation may be interpreted as deviation of modality synchronization from these implicitly chosen reference cell states. By convention, Δ*_i_* is computed with 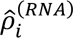 are reference. Therefore, Δ*_i_* > 0 represents states with higher density in ATAC and hence greater resolution in RNA. Conversely, Δ*_i_* < 0 represents states with greater resolution in ATAC.

#### Significance testing

The Bayesian posterior of Δ*_i_* has a variance:

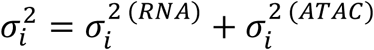

To assess the statistical significance of Δ*_i_*, we compute a posterior tail probability (PTP) that quantifies whether the observed density difference is larger than would be expected from posterior uncertainty alone, under the null hypothesis that the two modalities assign identical local density at state 𝑖 (Δ_true_ = 0). Given the Gaussian posterior on Δ*_i_*, the two-sided PTP is:

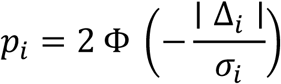

Where Φ is the standard normal cumulative distribution function. A state *i* is classified as significantly desynchronized when both

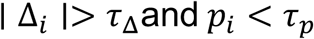

Where 𝜏_Δ_ and 𝜏*_p_* are user-specified thresholds on effect size and significance (defaults: 𝜏_Δ_ = log (2) and 𝜏*_p_* = 0.05). As Δᵢ, σᵢ, and the resulting PTP are defined at every cell in the dataset, Echo evaluates desynchronization at single-cell resolution without clustering or cell-type annotation.

### Echo Features

Desynchronized states between RNA and ATAC are a consequence of coordinated changes in expression of multiple genes, or in accessibility of multiple loci, without a commensurate change in the other modality. These changes are precisely what produces the resolution difference between modalities: the features driving the change track with state changes in the modality where they are active, while varying less across states defined by the other modality. As a result, when feature values are predicted from the cell-state space of each modality, desynchronized features are reconstructed more accurately in the higher-resolution modality than in the lower-resolution one.

Gene expression is used as features for identifying desynchronized genes in RNA-resolved states. Gene enhancer accessibility (**Methods**) and TF motif accessibility to identify desynchronized genes and TFs respectively, in ATAC-resolved states.

#### Per-modality feature prediction

Echo Features predicts the feature values using the modality-specific cell state space using Gaussian Process regression^21,22^. Specifically, we treat each feature value as a noisy measurement of an underlying state-driven signal and place a GP prior on that signal, yielding a posterior whose mean can be used as the prediction and whose variance quantifies uncertainty. Formally for a given feature with observed values 𝑦 = (𝑦_1_, …, 𝑦*_n_*) and a cell state embedding 𝑋^(K)^ where 𝑘 ∈ {𝑅𝑁𝐴, 𝐴𝑇𝐴𝐶}, we predict:

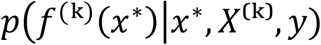

Where:

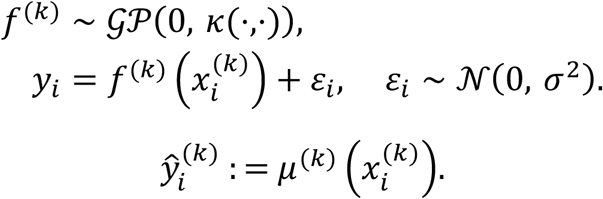

where 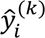 is the predicted value for cell 𝑖 for each modality 𝑘.

#### Difference in prediction accuracy

To assess the difference in prediction accuracy Echo calculates the Leave One Out (LOO) residuals as a per cell measure of prediction accuracy. LOO residuals are used to avoid underestimating the true error. The difference in prediction accuracy is computed for a group of related desynchronized states {𝑐}. This could be for example desynchronized states within a cell type or a cluster.

Concretely, for each cell 𝑖 and each modality 𝑘 we compute a leave-one-out predicted value 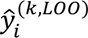 i.e., the GP posterior mean at cell 𝑖, fit without that cell. The LOO residual 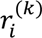 is computed as:

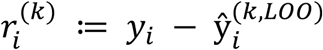

These per-cell LOO residuals are then aggregated within a cell group of interest (e.g. a cluster or cell type) 𝑐 to the per-modality mean-squared error 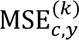 per feature 𝑦 as:

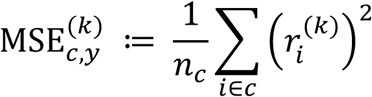

The difference in MSE between modalities,

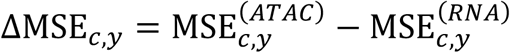

measures which cell-state space predicts the feature more accurately within the group.

Raw MSE differences are not directly comparable across features. In particular features with higher values (e.g. highly expressed genes) might have higher MSEs because proportionally small errors in prediction will lead to much higher MSEs. To address this Echo normalizes ΔMSE*_c,y_* by dividing them by the variance of the feature within the cell group 𝑐

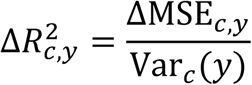

where Var*_c_*(𝑦) is the variance of the feature values in group *c*. This serves as a correction mechanism since features with high values often have high variances in both gene expression and chromatin accessibility due to overdispersion^70–72^. Further, dividing by the variance provides an intuitive link between 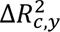 and the variance explained by each modality. Specifically,

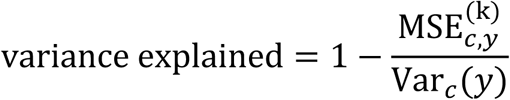

Thus, the quantity 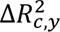 is equivalent to the difference between the variance of feature 𝑦 explained by the model trained on the first modality versus the variance of 𝑦 explained by the model trained on the second modality:

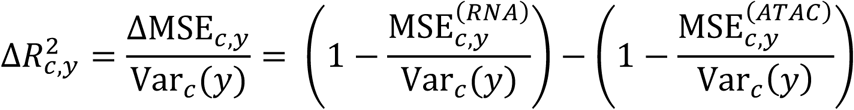

This approximately bounds values of 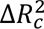 to [−1, 1]. By convention, positive values indicate features better predicted by RNA states and hence represents features underlying RNA-resolved states. Conversely, negative values indicate features better predicted by ATAC states and represent features underlying ATAC-resolved states.

#### Null distribution and significance testing

To assess the significance of 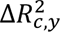, Echo compares it to null distribution generated by shuffling. For each feature, Echo randomly permutes that feature’s values across cells, breaking the connection between the feature and both cell-state.

The per-modality GP regression trained on the permuted values, yields null MSE differences and a null variance-explained difference 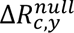 per feature and per group. The empirical mean and standard deviation of 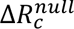 across features are used to computed Z-scores as:

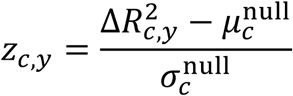

A two-sided p-value is derived from the standard normal distribution, and the resulting p-values are corrected within each group using Benjamini–Hochberg correction. A feature is classified as significantly desynchronized in group *c* with BH-adjusted p-value < 0.05.

Groups with fewer than a minimum number of cells (default: 50) are flagged and skipped to avoid unreliable MSE estimates.

#### Mahalanobis distance

As a complementary statistic, Echo also reports the Mahalanobis distance between the two per-modality prediction vectors. For a feature in group *c*, *R*^*n_c_*^ let be the vector of per-cell predictions 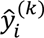 for 𝑖 ∈ 𝑐) from modality k, and let 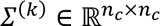 be the corresponding GP posterior. Unlike 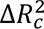, which compares within-modality variance-explained values, this is a direct cross-modality comparison of the predictions themselves. Within each group c we compute:

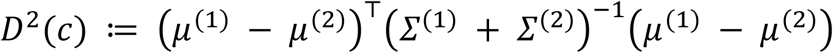

### Gene Enhancer Accessibility scores

Gene enhancer accessibility scores for each gene at single-cell resolution were computed by aggregating the accessibility of regulatory loci linked with the gene (**Supp. Fig. 4**). SEACells were first used to determine metacells using the ATAC modality and then to identify peaks correlated with each gene^73^. A weighted average of accessibility counts were computed for each peak and cell using the correlation as weights. The resulting weighted counts were normalized per cell and log transformed to derive gene enhancer accessibility scores.

## Code and Data Availability

Echo is available as an open-source codebase along with data at github.com/settylab/scEcho. A python package with documentation, tutorials and usage guidelines will be made available in a subsequent version.

## Author Contributions

C.F. and M.S. conceived and designed the study, developed Echo, developed additional analysis methods and statistical tests. D.J.O. provided statistical and interpretive input for conceptualization and implementation of Echo. C.F. performed all analysis. C.F., D.J.O., and M.S. wrote the manuscript.

## Acknowledgements

We thank Juliette Wohlschlegel, Matthew Wooten, Tom Reh and members of the Setty laboratory for discussions and comments on the manuscript. This study was supported by National Institute of General Medical Studies grant no. R35 GM147125 to M.S., Eunice Kennedy Shriver National Institute of Child Health and Human Development grant no. F31 HD118770 to C.F., National Institute of General Medical Studies grant no. K99GM159044 and the National Institutes of Health grant no. ORIP S10OD028685 to support high-performance computing at the Fred Hutchinson Cancer Research Center.

## References

1 Mitra, S. et al. Single-cell multi-ome regression models identify functional and disease-associated enhancers and enable chromatin potential analysis. Nat Genet 56, 627–636 (2024). 10.1038/s41588-024-01689-8

2 Hilliard, S. A. & El-Dahr, S. S. Epigenetics mechanisms in renal development. Pediatr Nephrol 31, 1055–1060 (2016). 10.1007/s00467-015-3228-x

3 Perino, M. & Veenstra, G. J. Chromatin Control of Developmental Dynamics and Plasticity. Dev Cell 38, 610–620 (2016). 10.1016/j.devcel.2016.08.004

4 Tao, L. et al. Enhancer decommissioning imposes an epigenetic barrier to sensory hair cell regeneration. Dev Cell 56, 2471–2485 e2475 (2021). 10.1016/j.devcel.2021.07.003

5 Kiani, K., Sanford, E. M., Goyal, Y. & Raj, A. Changes in chromatin accessibility are not concordant with transcriptional changes for single-factor perturbations. Mol Syst Biol 18, e10979 (2022). 10.15252/msb.202210979

6 Spitz, F. & Furlong, E. E. Transcription factors: from enhancer binding to developmental control. Nat Rev Genet 13, 613–626 (2012). 10.1038/nrg3207

7 Mercer, E. M. et al. Multilineage priming of enhancer repertoires precedes commitment to the B and myeloid cell lineages in hematopoietic progenitors. Immunity 35, 413–425 (2011). 10.1016/j.immuni.2011.06.013

8 Zaret, K. S. & Carroll, J. S. Pioneer transcription factors: establishing competence for gene expression. Genes Dev 25, 2227–2241 (2011). 10.1101/gad.176826.111

9 Levings, D. C., Lacher, S. E., Palacios-Moreno, J. & Slattery, M. Transcriptional reprogramming by oxidative stress occurs within a predefined chromatin accessibility landscape. Free Radic Biol Med 171, 319–331 (2021). 10.1016/j.freeradbiomed.2021.05.016

10 Bertoli, C., Skotheim, J. M. & de Bruin, R. A. Control of cell cycle transcription during G1 and S phases. Nat Rev Mol Cell Biol 14, 518–528 (2013). 10.1038/nrm3629

11 Ma, S. et al. Chromatin Potential Identified by Shared Single-Cell Profiling of RNA and Chromatin. Cell 183, 1103–1116 e1120 (2020). 10.1016/j.cell.2020.09.056

12 Cao, J. et al. Joint profiling of chromatin accessibility and gene expression in thousands of single cells. Science 361, 1380–1385 (2018). 10.1126/science.aau0730

13 Chen, S., Lake, B. B. & Zhang, K. High-throughput sequencing of the transcriptome and chromatin accessibility in the same cell. Nat Biotechnol 37, 1452–1457 (2019). 10.1038/s41587-019-0290-0

14 Li, C., Virgilio, M. C., Collins, K. L. & Welch, J. D. Multi-omic single-cell velocity models epigenome-transcriptome interactions and improves cell fate prediction. Nat Biotechnol 41, 387–398 (2023). 10.1038/s41587-022-01476-y

15 Fu, S. et al. Benchmarking single-cell multi-modal data integrations. Nat Methods 22, 2437–2448 (2025). 10.1038/s41592-025-02737-9

16 Cao, Z. J. & Gao, G. Multi-omics single-cell data integration and regulatory inference with graph-linked embedding. Nat Biotechnol 40, 1458–1466 (2022). 10.1038/s41587-022-01284-4

17 Lin, K. Z. & Zhang, N. R. Quantifying common and distinct information in single-cell multimodal data with Tilted Canonical Correlation Analysis. Proc Natl Acad Sci U S A 120, e2303647120 (2023). 10.1073/pnas.2303647120

18 Argelaguet, R. et al. MOFA+: a statistical framework for comprehensive integration of multi-modal single-cell data. Genome Biol 21, 111 (2020). 10.1186/s13059-020-02015-1

19 Hao, Y. et al. Integrated analysis of multimodal single-cell data. Cell 184, 3573–3587 e3529 (2021). 10.1016/j.cell.2021.04.048

20 Wang, F. A. et al. scTFBridge: a disentangled deep generative model informed by TF-motif binding for gene regulation inference in single-cell multi-omics. Nat Commun 16, 9166 (2025). 10.1038/s41467-025-64227-y

21 Otto, D. J., Jordan, C., Dury, B., Dien, C. & Setty, M. Quantifying cell-state densities in single-cell phenotypic landscapes using Mellon. Nat Methods 21, 1185–1195 (2024). 10.1038/s41592-024-02302-w

22 Seeger, M. Gaussian processes for machine learning. Int J Neural Syst 14, 69–106 (2004). 10.1142/S0129065704001899

23 Wohlschlegel, J. et al. ASCL1 induces neurogenesis in human Muller glia. Stem Cell Reports 18, 2400–2417 (2023). 10.1016/j.stemcr.2023.10.021

24 Diacou, R. et al. Cell fate decisions, transcription factors and signaling during early retinal development. Prog Retin Eye Res 91, 101093 (2022). 10.1016/j.preteyeres.2022.101093

25 Lyu, P. et al. Gene regulatory networks controlling temporal patterning, neurogenesis, and cell-fate specification in mammalian retina. Cell Rep 37, 109994 (2021). 10.1016/j.celrep.2021.109994

26 Thomas, E. D. et al. Cell-specific cis-regulatory elements and mechanisms of non-coding genetic disease in human retina and retinal organoids. Dev Cell 57, 820–836 e826 (2022). 10.1016/j.devcel.2022.02.018

27 Finkbeiner, C. et al. Single-cell ATAC-seq of fetal human retina and stem-cell-derived retinal organoids shows changing chromatin landscapes during cell fate acquisition. Cell Rep 38, 110294 (2022). 10.1016/j.celrep.2021.110294

28 Turner, D. L. & Cepko, C. L. A common progenitor for neurons and glia persists in rat retina late in development. Nature 328, 131–136 (1987). 10.1038/328131a0

29 Turner, D. L., Snyder, E. Y. & Cepko, C. L. Lineage-independent determination of cell type in the embryonic mouse retina. Neuron 4, 833–845 (1990). 10.1016/0896-6273(90)90136-4

30 Godinho, L. et al. Nonapical symmetric divisions underlie horizontal cell layer formation in the developing retina in vivo. Neuron 56, 597–603 (2007). 10.1016/j.neuron.2007.09.036

31 Hafler, B. P. et al. Transcription factor Olig2 defines subpopulations of retinal progenitor cells biased toward specific cell fates. Proc Natl Acad Sci U S A 109, 7882–7887 (2012). 10.1073/pnas.1203138109

32 Cepko, C. Intrinsically different retinal progenitor cells produce specific types of progeny. Nat Rev Neurosci 15, 615–627 (2014). 10.1038/nrn3767

33 Suzuki, S. C. et al. Cone photoreceptor types in zebrafish are generated by symmetric terminal divisions of dedicated precursors. Proc Natl Acad Sci U S A 110, 15109–15114 (2013). 10.1073/pnas.1303551110

34 Capecchi, M. R. & Pozner, A. ASPM regulates symmetric stem cell division by tuning Cyclin E ubiquitination. Nat Commun 6, 8763 (2015). 10.1038/ncomms9763

35 Heck, M. M., Hittelman, W. N. & Earnshaw, W. C. Differential expression of DNA topoisomerases I and II during the eukaryotic cell cycle. Proc Natl Acad Sci U S A 85, 1086–1090 (1988). 10.1073/pnas.85.4.1086

36 Zhu, X. et al. Characterization of a novel 350-kilodalton nuclear phosphoprotein that is specifically involved in mitotic-phase progression. Mol Cell Biol 15, 5017–5029 (1995). 10.1128/MCB.15.9.5017

37 Tirosh, I. et al. Dissecting the multicellular ecosystem of metastatic melanoma by single-cell RNA-seq. Science 352, 189–196 (2016). 10.1126/science.aad0501

38 Muller, G. A. & Engeland, K. The central role of CDE/CHR promoter elements in the regulation of cell cycle-dependent gene transcription. FEBS J 277, 877–893 (2010). 10.1111/j.1742-4658.2009.07508.x

39 Sadasivam, S. & DeCaprio, J. A. The DREAM complex: master coordinator of cell cycle-dependent gene expression. Nat Rev Cancer 13, 585–595 (2013). 10.1038/nrc3556

40 Ly, L. L., Yoshida, H. & Yamaguchi, M. Nuclear transcription factor Y and its roles in cellular processes related to human disease. Am J Cancer Res 3, 339–346 (2013).

41 Nishida, A. et al. Otx2 homeobox gene controls retinal photoreceptor cell fate and pineal gland development. Nat Neurosci 6, 1255–1263 (2003). 10.1038/nn1155

42 Sato, S. et al. Dkk3-Cre BAC transgenic mouse line: a tool for highly efficient gene deletion in retinal progenitor cells. Genesis 45, 502–507 (2007). 10.1002/dvg.20318

43 Yamamoto, H., Kon, T., Omori, Y. & Furukawa, T. Functional and Evolutionary Diversification of Otx2 and Crx in Vertebrate Retinal Photoreceptor and Bipolar Cell Development. Cell Rep 30, 658–671 e655 (2020). 10.1016/j.celrep.2019.12.072

44 Ghinia Tegla, M. G., et al. OTX2 represses sister cell fate choices in the developing retina to promote photoreceptor specification. Elife 9 (2020). 10.7554/eLife.54279

45 Katoh, K. et al. Blimp1 suppresses Chx10 expression in differentiating retinal photoreceptor precursors to ensure proper photoreceptor development. J Neurosci 30, 6515–6526 (2010). 10.1523/JNEUROSCI.0771-10.2010

46 Brzezinski, J. A. t., Lamba, D. A. & Reh, T. A. Blimp1 controls photoreceptor versus bipolar cell fate choice during retinal development. Development 137, 619–629 (2010). 10.1242/dev.043968

47 Setty, M. et al. Characterization of cell fate probabilities in single-cell data with Palantir. Nat Biotechnol 37, 451–460 (2019). 10.1038/s41587-019-0068-4

48 Kaufman, M. L. et al. Initiation of Otx2 expression in the developing mouse retina requires a unique enhancer and either Ascl1 or Neurog2 activity. Development 148 (2021). 10.1242/dev.199399

49 Henikoff, S. et al. RNA polymerase II at histone genes predicts outcome in human cancer. Science 387, 737–743 (2025). 10.1126/science.ads2169

50 Imayoshi, I. & Kageyama, R. bHLH factors in self-renewal, multipotency, and fate choice of neural progenitor cells. Neuron 82, 9–23 (2014). 10.1016/j.neuron.2014.03.018

51 Baker, N. E. & Brown, N. L. All in the family: proneural bHLH genes and neuronal diversity. Development 145 (2018). 10.1242/dev.159426

52 Maurer, K. A., Riesenberg, A. N. & Brown, N. L. Notch signaling differentially regulates Atoh7 and Neurog2 in the distal mouse retina. Development 141, 3243–3254 (2014). 10.1242/dev.106245

53 Chen, S. et al. Crx, a novel Otx-like paired-homeodomain protein, binds to and transactivates photoreceptor cell-specific genes. Neuron 19, 1017–1030 (1997). 10.1016/s0896-6273(00)80394-3

54 Hennig, A. K., Peng, G. H. & Chen, S. Regulation of photoreceptor gene expression by Crx-associated transcription factor network. Brain Res 1192, 114–133 (2008). 10.1016/j.brainres.2007.06.036

55 Leigh, A., Swaroop, A., Kruczek, K., Ullah, E. & Brooks, B. P. Cone Rod Homeobox (CRX): literature review and new insights. Ophthalmic Genet 46, 338–346 (2025). 10.1080/13816810.2025.2458086

56 Gan, L. et al. POU domain factor Brn-3b is required for the development of a large set of retinal ganglion cells. Proc Natl Acad Sci U S A 93, 3920–3925 (1996). 10.1073/pnas.93.9.3920

57 Ivanov, D. Notch Signaling-Induced Oscillatory Gene Expression May Drive Neurogenesis in the Developing Retina. Front Mol Neurosci 12, 226 (2019). 10.3389/fnmol.2019.00226

58 Shimojo, H., Ohtsuka, T. & Kageyama, R. Oscillations in notch signaling regulate maintenance of neural progenitors. Neuron 58, 52–64 (2008). 10.1016/j.neuron.2008.02.014

59 Persad, S. et al. SEACells infers transcriptional and epigenomic cellular states from single-cell genomics data. Nat Biotechnol (2023). 10.1038/s41587-023-01716-9

60 Zon, L. I. Intrinsic and extrinsic control of haematopoietic stem-cell self-renewal. Nature 453, 306–313 (2008). 10.1038/nature07038

61 Orkin, S. H. Diversification of haematopoietic stem cells to specific lineages. Nat Rev Genet 1, 57–64 (2000). 10.1038/35049577

62 Rosenbauer, F. & Tenen, D. G. Transcription factors in myeloid development: balancing differentiation with transformation. Nat Rev Immunol 7, 105–117 (2007). 10.1038/nri2024

63 Okada, S., Fukuda, T., Inada, K. & Tokuhisa, T. Prolonged expression of c-fos suppresses cell cycle entry of dormant hematopoietic stem cells. Blood 93, 816–825 (1999).

64 Chen, J. et al. Fosl2 Deficiency Predisposes Mice to Osteopetrosis, Leading to Bone Marrow Failure. J Immunol 212, 1081–1093 (2024). 10.4049/jimmunol.2300592

65 Andrews, N. C., Erdjument-Bromage, H., Davidson, M. B., Tempst, P. & Orkin, S. H. Erythroid transcription factor NF-E2 is a haematopoietic-specific basic-leucine zipper protein. Nature 362, 722–728 (1993). 10.1038/362722a0

66 Murre, C. ’Big bang’ of B-cell development revealed. Genes Dev 32, 93–95 (2018). 10.1101/gad.311357.118

67 May, G. et al. Dynamic analysis of gene expression and genome-wide transcription factor binding during lineage specification of multipotent progenitors. Cell Stem Cell 13, 754–768 (2013). 10.1016/j.stem.2013.09.003

68 Coifman, R. R. et al. Geometric diffusions as a tool for harmonic analysis and structure definition of data: diffusion maps. Proc Natl Acad Sci U S A 102, 7426–7431 (2005). 10.1073/pnas.0500334102

69 Haghverdi, L., Buttner, M., Wolf, F. A., Buettner, F. & Theis, F. J. Diffusion pseudotime robustly reconstructs lineage branching. Nat Methods 13, 845–848 (2016). 10.1038/nmeth.3971

70 Ahlmann-Eltze, C. & Huber, W. Comparison of transformations for single-cell RNA-seq data. Nat Methods 20, 665–672 (2023). 10.1038/s41592-023-01814-1

71 Hafemeister, C. & Satija, R. Normalization and variance stabilization of single-cell RNA-seq data using regularized negative binomial regression. Genome Biol 20, 296 (2019). 10.1186/s13059-019-1874-1

72 Das, S. & Rai, S. N. Statistical methods for analysis of single-cell RNA-sequencing data. MethodsX 8, 101580 (2021). 10.1016/j.mex.2021.101580

73 Persad, S. et al. SEACells infers transcriptional and epigenomic cellular states from single-cell genomics data. Nat Biotechnol 41, 1746–1757 (2023). 10.1038/s41587-023-01716-9

74 Schep, A. N., Wu, B., Buenrostro, J. D. & Greenleaf, W. J. chromVAR: inferring transcription-factor-associated accessibility from single-cell epigenomic data. Nat Methods 14, 975–978 (2017). 10.1038/nmeth.4401

