## Supplementary Figures for "Echo characterizes the desynchronization of gene expression and chromatin accessibility during cell-state transitions"

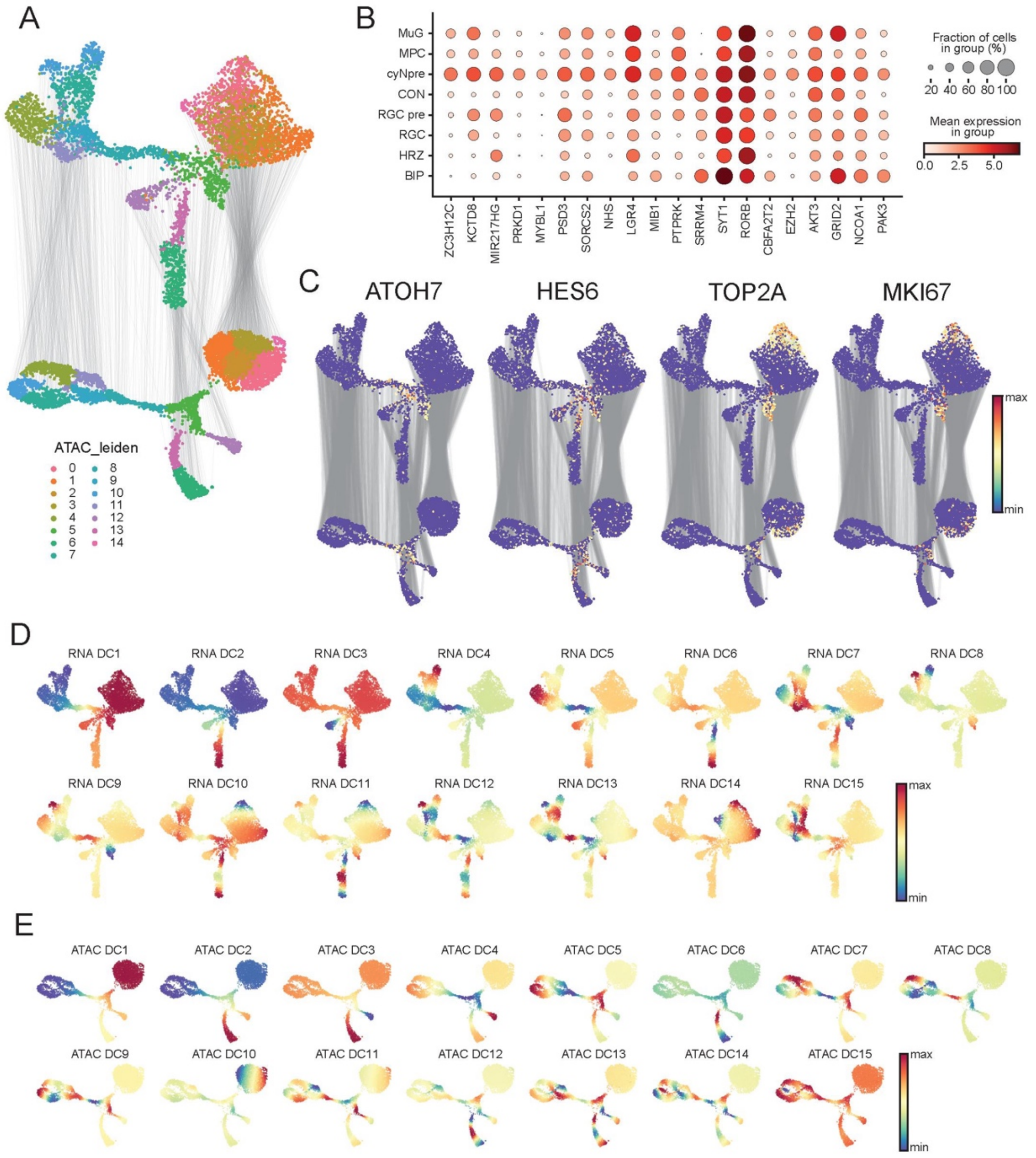

Supplementary Figure 1: Paired single-cell RNA- and ATAC-seq data of fetal retina development

(A) UMAPs from **Fig. 2A**, colored by ATAC leiden clusters. ATAC clusters were used to annotate cycling neurogenic precursors (cyNpre cells).

(B) Dot plot showing the expression of top 20 differentially expressed genes in cyNpre cells.

(C) UMAPs from 2A, colored by gene expression of ATOH7, HES6, and cycling cell markers. These markers informed the cyNpre annotation

(D, E) RNA UMAPs (D) and ATAC UMAPs (E) colored by respective diffusion components. These are the state representations used as input to Echo.

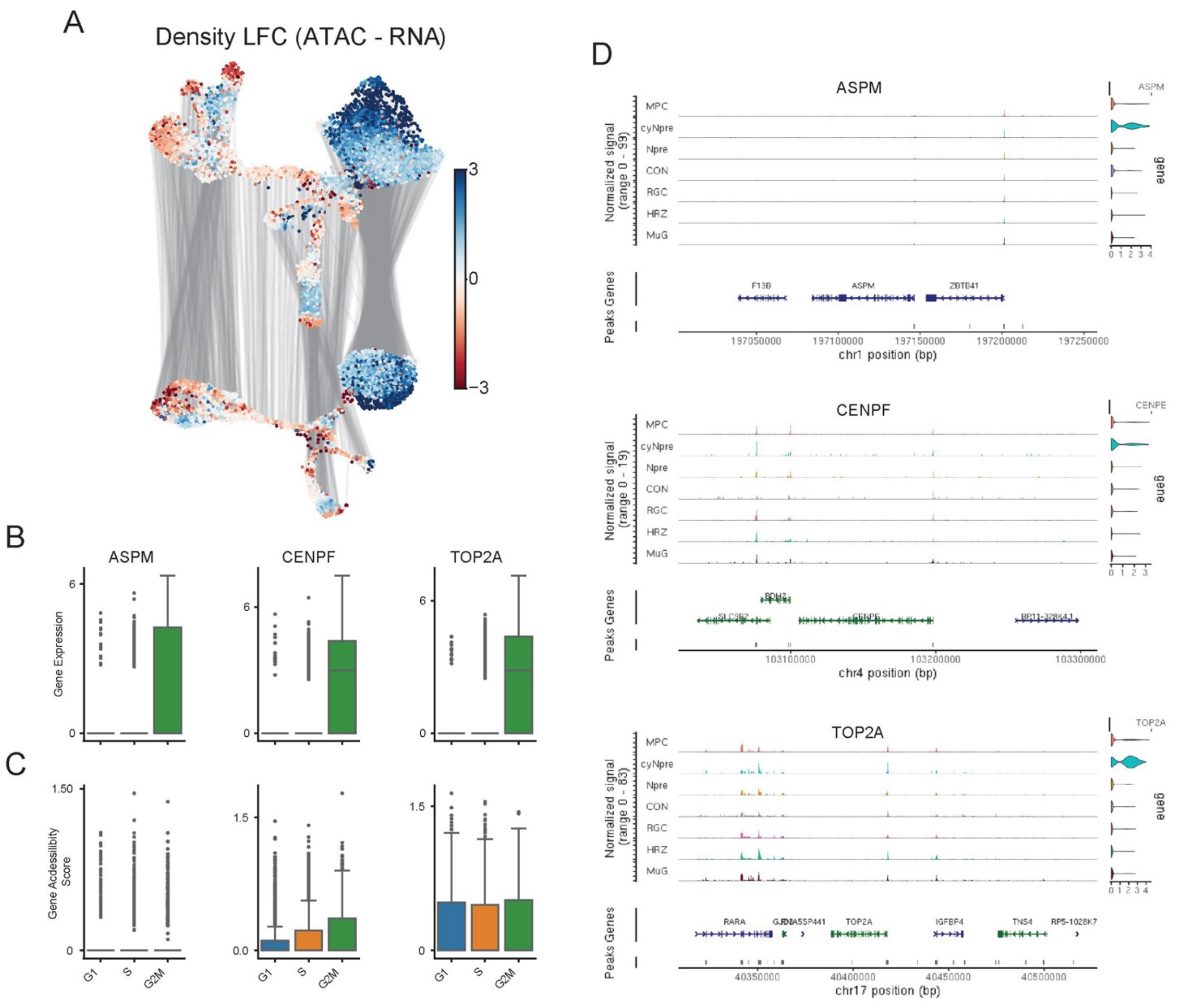

### Supplementary Figure 2: MPCs are resolved better in RNA than ATAC due to cell cycle

(A) UMAPs from **Fig. 2A**, colored by density log fold change. Positive fold change indicates greater resolution in RNA and negative fold change indicates greater resolution in ATAC.

(B) Boxplots showing expression of the top desynchronized genes from **Fig. 2E**, in G1, S and G2M cell cycle phase MPCs.

(C) Same as (B) using gene accessibility scores from ARCHR[1].

(D) Coverage plots for genes in (B)

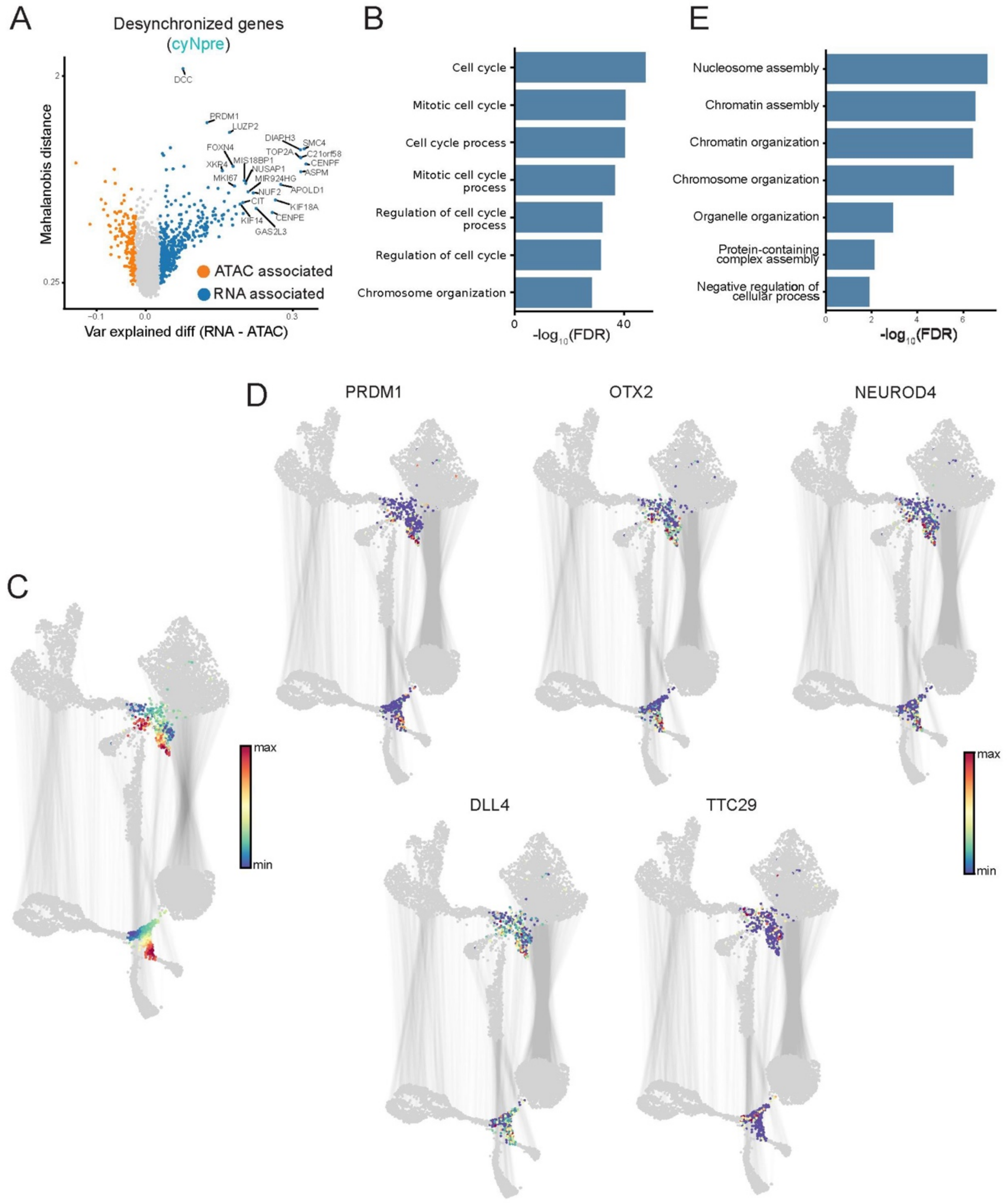

### Supplementary Figure 3: Desynchronization of cyNpre cells

(A) Echo Features applied to gene expression in cyNpre cells. Pseudo-volcano plot shows difference in variance explained (RNA − ATAC) on the x-axis and Mahalanobis distance on the y-axis. Significantly desynchronized genes are colored by their modality association, and top-ranked genes are labeled.

(B) Gene ontology analysis of RNA associated genes from (A)**.**

(C) UMAPs from **Fig. 2A**, colored by ATAC diffusion component 9 that illustrates the cyNpre state organization in ATAC. Non-cyNpre cells are in grey.

(D) Same as (B), colored by top 5 desynchronized genes based on enhancer accessibility from **Fig. 3B**.

(E) Gene ontology analysis of RNA associated genes from **Fig. 3B**.

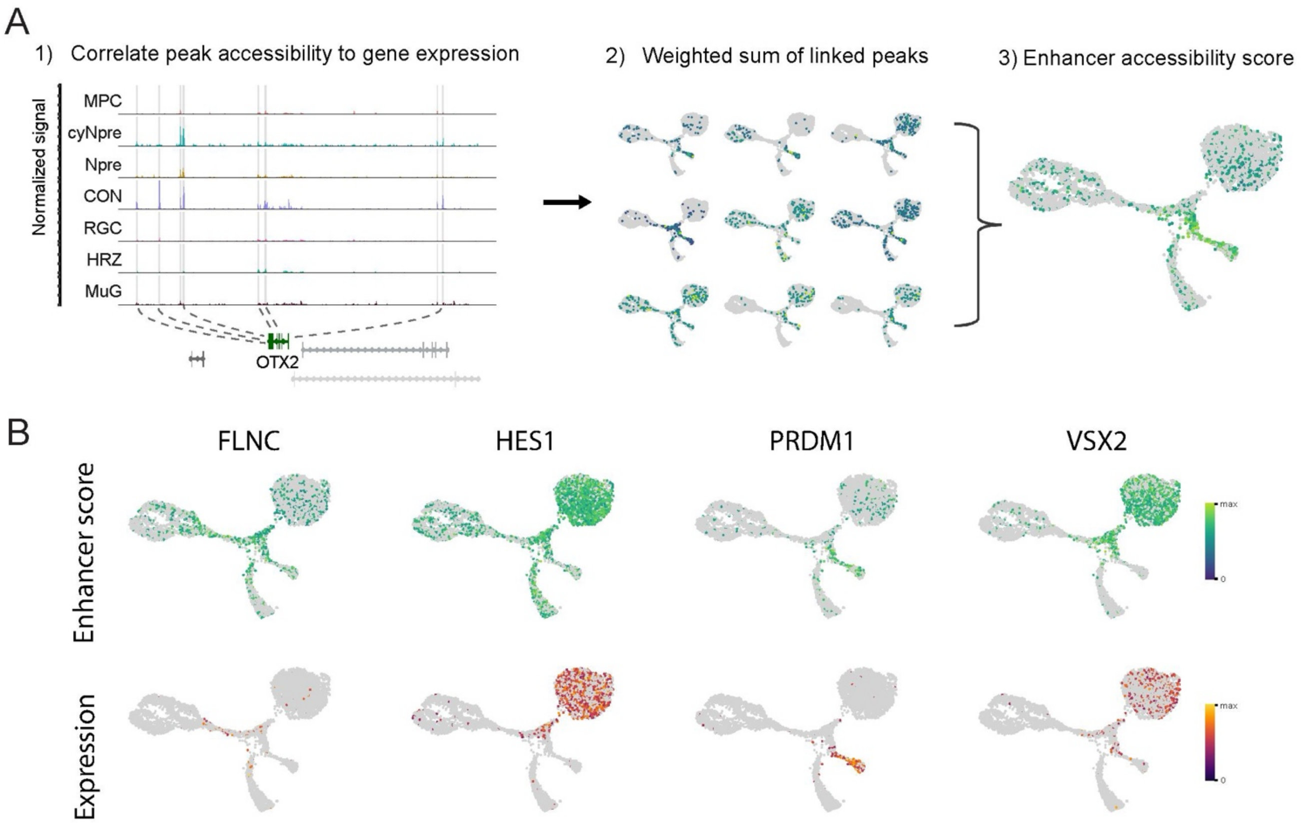

### Supplementary Figure 4: Computation of enhancer accessibility scores

(A) Illustration of enhancer accessibility score for each gene using OTX2 as an example. 1. Coverage plots highlighting the set of peaks with accessibility correlated with OTX2 expression. SEACells metacells[2] are used for computing correlations. 2. Correlation-weighted average of peak accessibility per cell is used as the enhancer accessibility score for the gene (3).

(B) ATAC UMAPs from **Fig. 2A**, colored by enhancer scores (top) and gene expression (bottom) for a selection of genes with desynchronized enhancer scores from **Fig. 3B**.

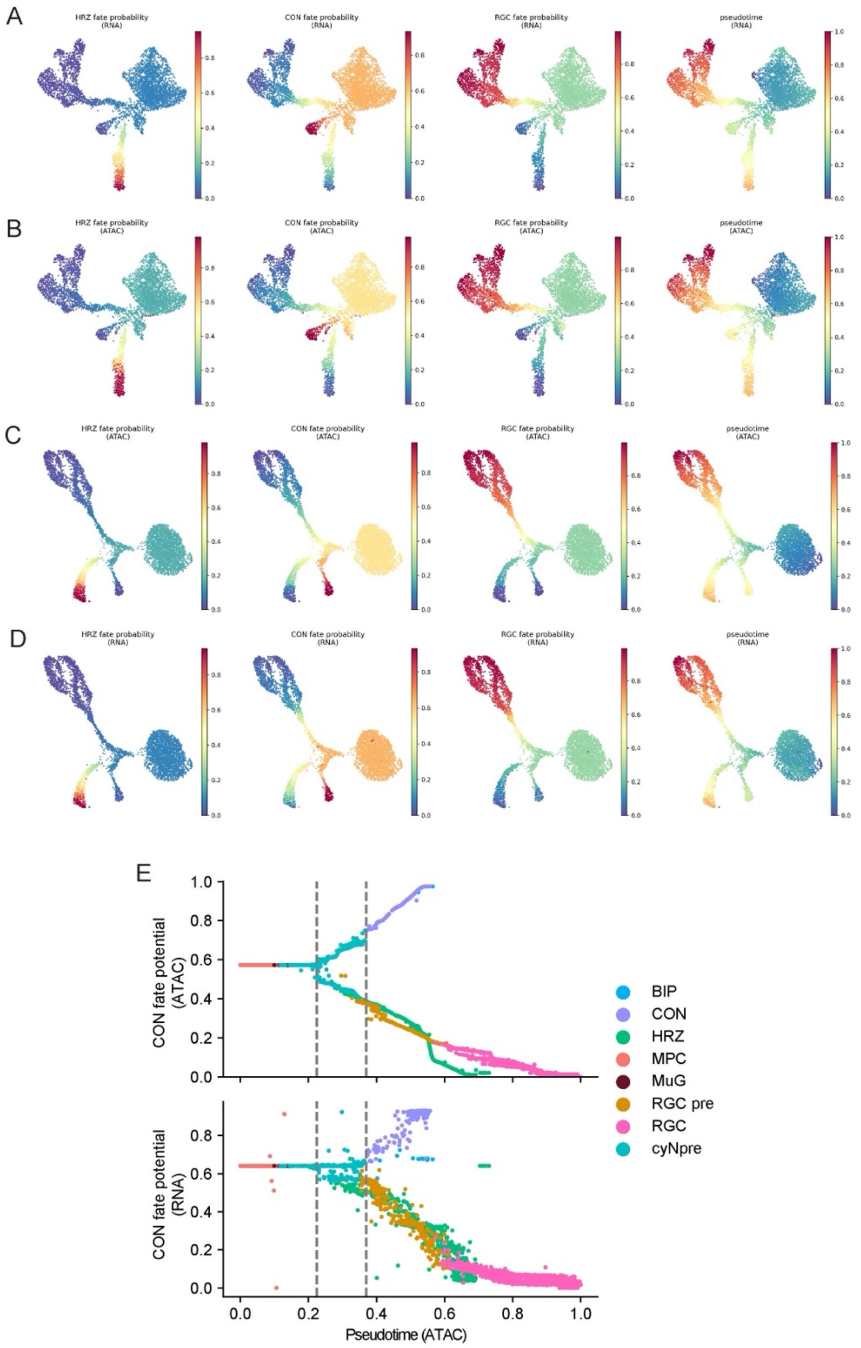

### Supplementary Figure 5: Fate probabilities in retinal development

(A) RNA UMAPs from **Fig 2A.**, colored by fate propensities and pseudotime computed from RNA using Palantir[3].

(B) RNA UMAPs colored by fate propensities and pseudotime computed from ATAC.

(C) Same as (A), colored on ATAC UMAPs.

(D) Same as (B), colored on ATAC UMAPs.

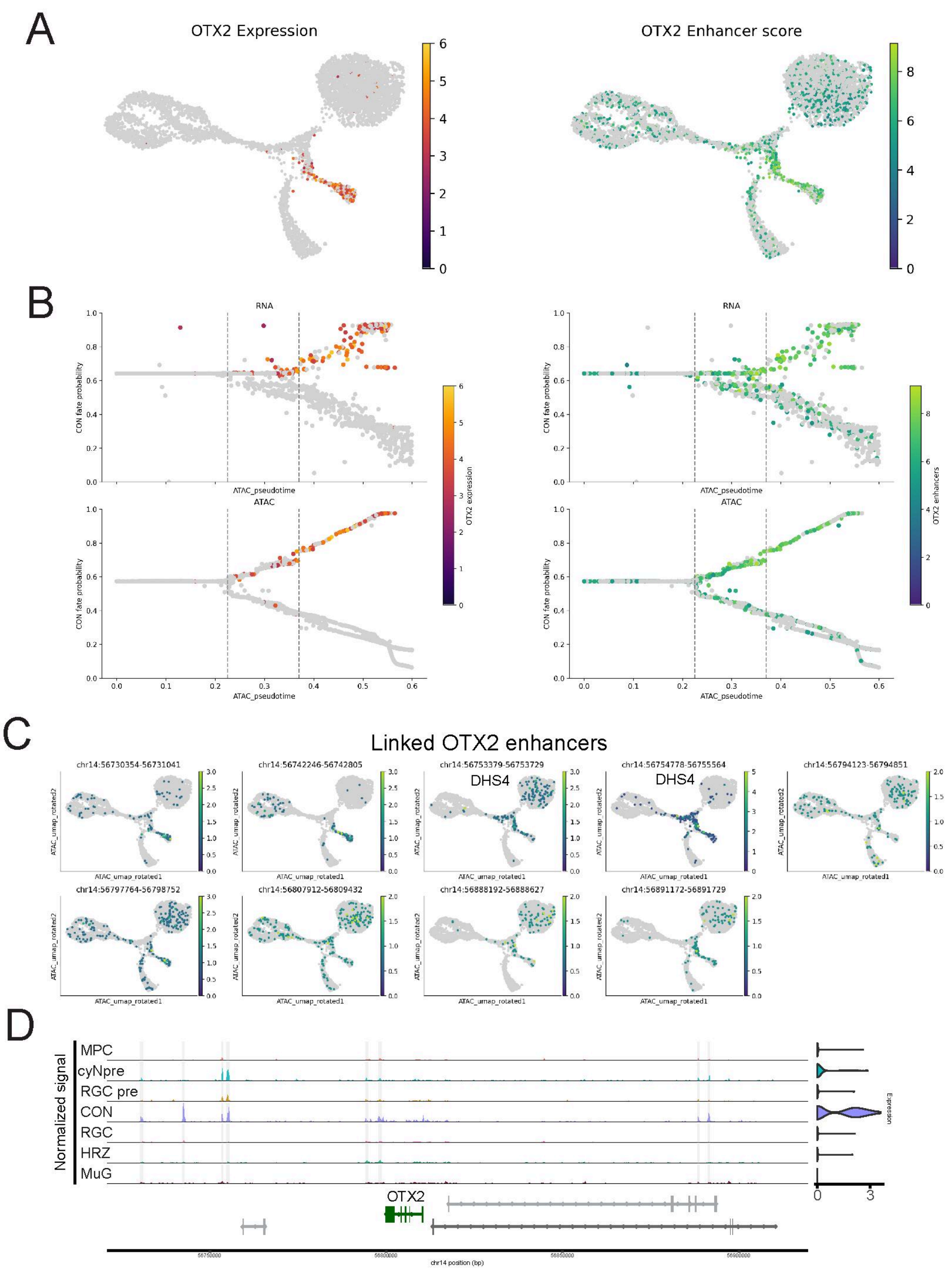

### Supplementary Figure 6: Interaction of OTX2 gene expression and chromatin accessibility

(A) ATAC UMAP from **Fig. 2A**, colored by OTX2 expression (left) and OTX2 enhancer accessibility score (right)

(B) Same as **Fig. 3E**, colored by OTX2 expression (left) and OTX2 enhancer accessibility score (right).

(C) ATAC UMAP from **Fig. 2A**, colored by accessibility of enhancers linked to OTX2. Enhancer accessibility scores in (A) and (B) are weighted average of these enhancers.

(D) Right: Coverage plots of the OTX2 locus across major cell types. Left: OTX2 expression in the same cells.

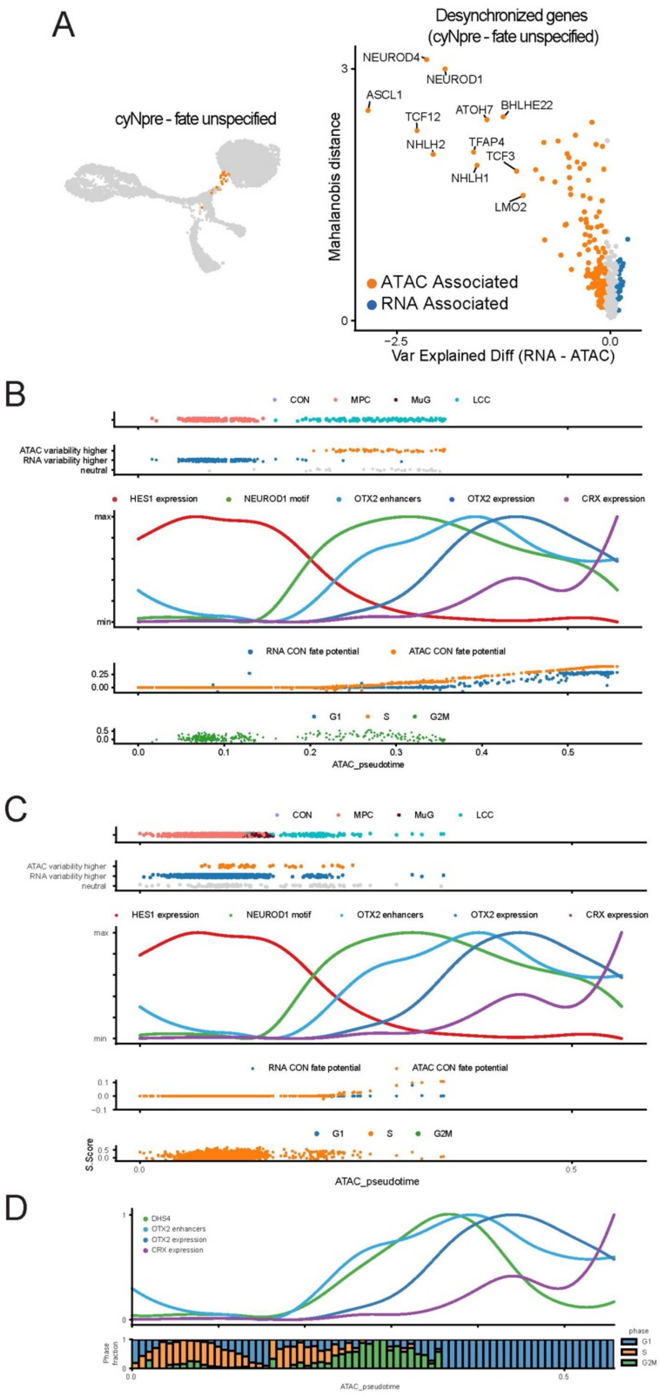

Supplementary Figure 7: Desynchronization reveals that exit of multipotency and priming of cone fate is coupled through cell cycle.

(A) Right: ATAC UMAP from **Fig. 2A**, highlighted by ATAC resolved cells in the MPC-cyNpre transition **Fig. 2C**. Left: Echo Features applied to gene enhancer accessibility scores in these cells. Pseudo-volcano plot shows difference in variance explained (RNA − ATAC) on the x-axis and Mahalanobis distance on the y-axis. Significantly desynchronized genes are colored by their modality association, and top-ranked genes are labeled.

(B) Same as **Fig. 3J**, showing cells annotated as in G2M phase of the cell cycle only. Dynamics in the third panel from the top are same as **Fig. 3J**.

(C) Same as (B) showing cells annotated as in S phase of the cell cycle only.

(D) Dynamics of OTX2 expression, enhancer accessibility score, and OTX2 linked DHS4 enhancer accessibility along ATAC pseudotime.

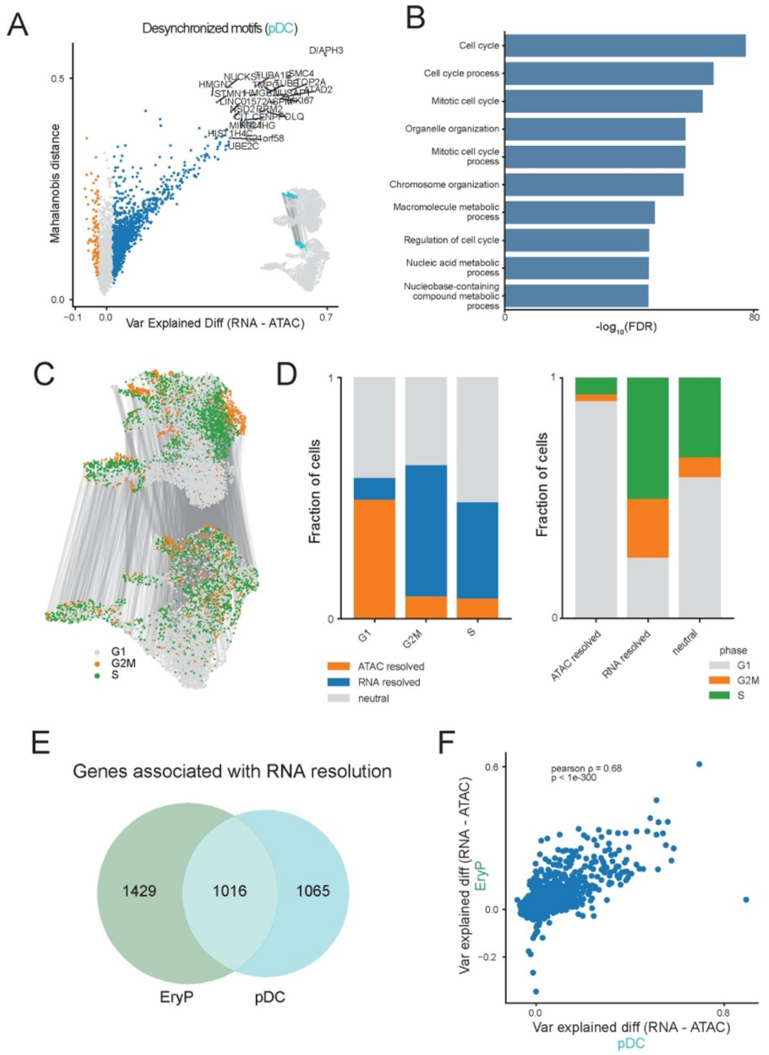

### Supplementary Figure 8: RNA resolution is associated with cell cycle in hematopoietic differentiation

(A) Echo Features applied to gene expression in the RNA-resolved pDC states. Pseudo-volcano plot shows difference in variance explained (RNA − ATAC) on the x-axis and Mahalanobis distance as the significance metric on the y-axis. Significantly desynchronized genes are colored by their modality association, and top-ranked genes are labeled. pDCs are highlighted in the inset on UMAPs from **Fig. 4A**.

(B) Gene Ontology terms enriched in the pDC desynchronized gene set

(C) UMAPs from **Fig. 4A**, colored by cell cycle phase.

(D) Left: Desynchronized states across cell cycle phase in hematopoietic differentiation. Right: Cell cycle phase of desynchronized states.

(E) Venn diagram of genes association with RNA resolution between erythroid precursors and pDCs.

(F) Correlation of Echo Features metrics between pDCs and erythroid precursors.

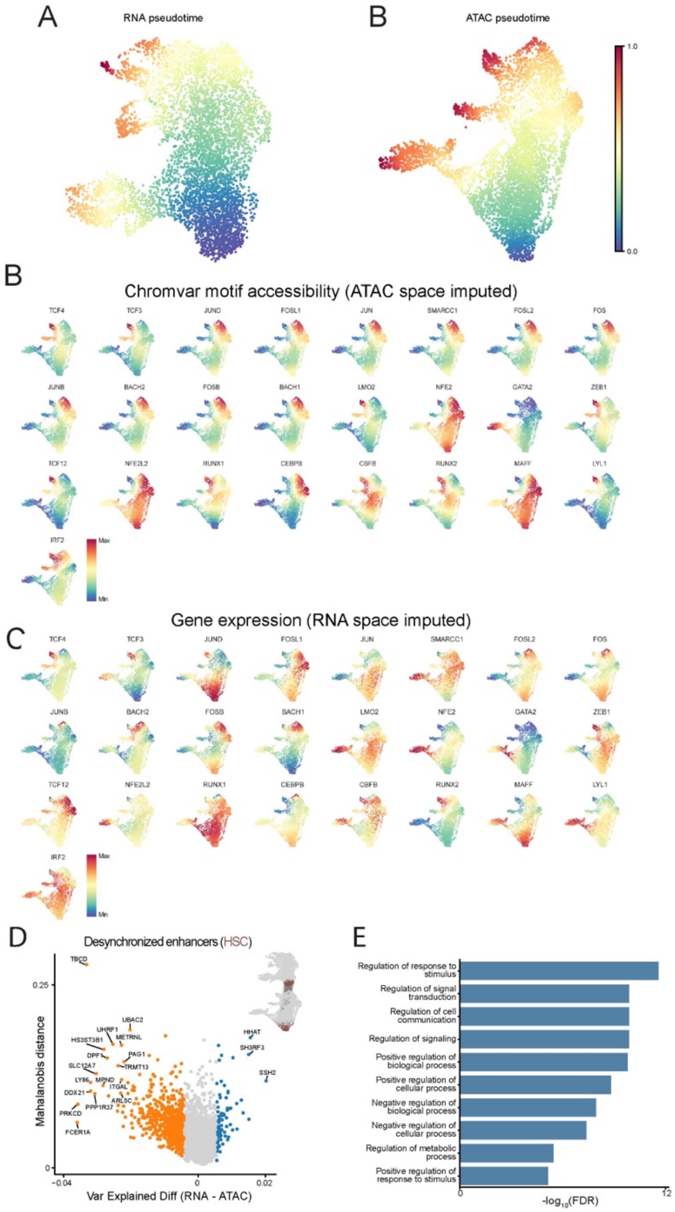

### Supplementary Figure 9: Balance between HSC quiescence and differentiation is better resolved by ATAC than RNA

(A) Left: RNA UMAP from **Fig. 4A**, colored by RNA pseudotime. Right: ATAC UMAP from **Fig. 4A**, colored by ATAC pseudotime.

(B) ATAC UMAP from **Fig. 4A**, colored by imputed chromVAR scores for TFs in **Fig. 4E**.

(C) Same as (B), colored by imputed gene expression.

(D) Echo Features applied to gene enhancer accessibility scores in HSCs Pseudo-volcano plot shows difference in variance explained (RNA − ATAC) on the x-axis and Mahalanobis distance on the y-axis. Significantly desynchronized genes are colored by their modality association, and top-ranked genes are labeled.

(E) Gene ontology analysis of desynchronized genes from (D).

### References

1 Granja, J. M. *et al.* ArchR is a scalable software package for integrative single-cell chromatin accessibility analysis. *Nat Genet* **53**, 403-411 (2021). https://doi.org/10.1038/s41588-021-00790-6

2 Persad, S. *et al.* SEACells infers transcriptional and epigenomic cellular states from single-cell genomics data. *Nat Biotechnol* **41**, 1746-1757 (2023). https://doi.org/10.1038/s41587-023-01716-9

3 Setty, M. *et al.* Characterization of cell fate probabilities in single-cell data with Palantir. *Nat Biotechnol* **37**, 451-460 (2019). https://doi.org/10.1038/s41587-019-0068-4
